# Genomically Complex Human Angiosarcoma and Canine Hemangiosarcoma Establish Convergent Angiogenic Transcriptional Programs Driven by Novel Gene Fusions

**DOI:** 10.1101/2020.08.11.246777

**Authors:** Jong Hyuk Kim, Kate Megquier, Rachael Thomas, Aaron L. Sarver, Jung Min Song, Yoon Tae Kim, Nuojin Cheng, Ashley J. Schulte, Michael A. Linden, Paari Murugan, LeAnn Oseth, Colleen L. Forster, Ingegerd Elvers, Ross Swofford, Jason Turner-Maier, Elinor K. Karlsson, Matthew Breen, Kerstin Lindblad-Toh, Jaime F. Modiano

**Affiliations:** Animal Cancer Care and Research Program, University of Minnesota, St Paul, MN, USA; Department of Veterinary Clinical Sciences, College of Veterinary Medicine, University of Minnesota, St Paul, MN, USA; Masonic Cancer Center, University of Minnesota, Minneapolis, MN, USA; Institute for Engineering in Medicine, University of Minnesota, Minneapolis, MN, USA; Broad Institute of Harvard and MIT, Cambridge, MA, USA; Department of Molecular Biomedical Sciences, College of Veterinary Medicine & Comparative Medicine Institute, North Carolina State University, Raleigh, NC, USA; Institute for Health Informatics, University of Minnesota, Minneapolis, MN, USA; Department of Electrical Engineering and Computer Science, York University, Toronto, Ontario, Canada; School of Mathematics, College of Science and Engineering, University of Minnesota, Minneapolis, MN, USA; Department of Laboratory Medicine and Pathology, School of Medicine, University of Minnesota, Minneapolis, MN, USA; The University of Minnesota Biological Materials Procurement Network (BioNet), University of Minnesota, Minneapolis, MN, USA; Science for Life Laboratory, Department of Medical Biochemistry and Microbiology, Uppsala University, Uppsala, Sweden; University of Massachusetts Medical School, Worcester, MA, USA; Cancer Genetics Program, University of North Carolina Lineberger Comprehensive Cancer Center, Raleigh, NC, USA; Center for Immunology, University of Minnesota, Minneapolis, MN, USA; Stem Cell Institute, University of Minnesota, Minneapolis, MN, USA

**Keywords:** Angiosarcoma, chromosome translocation, fusion gene, hemangiosarcoma, *TP53*

## Abstract

Sporadic angiosarcomas (ASs) are aggressive vascular sarcomas whose rarity and genomic complexity present significant obstacles in deciphering the pathogenic significance of individual genetic alterations. Numerous fusion genes have been identified across multiple types of cancers, but their existence and significance remain unclear in sporadic ASs. In this study, we leveraged RNA sequencing data from thirteen human ASs and 76 spontaneous canine hemangiosarcomas (HSAs) to identify fusion genes associated with spontaneous vascular malignancies. Ten novel protein-coding fusion genes, including *TEX2-PECAM1* and *ATP8A2-FLT1,* were identified in seven of the thirteen human tumors, with two tumors showing mutations of *TP53. HRAS* and *NRAS* mutations were found in ASs without fusions or *TP53* mutations. We found fifteen novel protein-coding fusion genes including *MYO16-PTK2, GABRA3-FLT1,* and *AKT3-XPNPEP1* in eleven of the 76 canine HSAs; these fusion genes were seen exclusively in tumors of the angiogenic molecular subtype that contained recurrent mutations in *TP53, PIK3CA, PIK3R1,* and *NRAS.* In particular, fusion genes and mutations of *TP53* co-occurred in tumors with higher frequency than expected by random chance, and they enriched gene signatures predicting activation of angiogenic pathways. Comparative transcriptomic analysis of human ASs and canine HSAs identified shared molecular signatures associated with activation of PI3K/AKT/mTOR pathways. Our data show that, while driver events of malignant vasoformative tumors of humans and dogs include diverse mutations and stochastic rearrangements that create novel fusion genes, convergent transcriptional programs govern the highly conserved morphological organization and biological behavior of these tumors in both species.

## Introduction

Sarcomas are diverse tumors that arise from cells of mesenchymal origin in soft tissues such as blood and lymphatic vessels, fat, bone, cartilage, muscle, and connective tissues. The heterogeneity of sarcomas has provided an impetus for developing molecular approaches to classify these tumors (1,2), leading to their categorization into genomically simple and genomically complex sarcomas (1,3). Angiosarcomas (ASs) are rare, highly aggressive, genomically complex sarcomas of blood vessel-forming cells (3,4). The five-year survival rate of AS is approximately 40% (5-7), but half of patients have metastatic or unresectable disease with a median overall survival of less than 6 months (8). The events that drive progression are incompletely understood; previous studies have identified recurrent mutations of *RAS, PTPRB, PLCG1, KDR* (kinase insert domain receptor, also known as *VEGFR2), TP53, PIK3CA,* and *FLT4 (VEGFR3)* in human ASs (9-12). *MYC* gene amplification and alterations in the TP53, CDKN2, NF-κB/IL-6, PIK3CA/AKT/mTOR pathways have also been reported (13); however, these studies represent a small case series, precluding definitive conclusions regarding pathogenic mechanisms that contribute to the genetic cause and to the progression of the disease.

Hemangiosarcoma (HSA) is a malignant vascular tumor that is common in dogs with an estimated tens of thousands of cases diagnosed each year (14-16). Canine HSA shares clinical and morphological features with human AS, as well as aspects of its mutational landscape (17-20). We previously documented three molecular subtypes of HSA, characterized by angiogenic, inflammatory, and adipogenic transcriptomic signatures (21). These gene expression signatures are conserved in HSA progenitor cells that show multipotency and self-renewal (21). Nevertheless, the transcriptional state of these HSA progenitor cells seems to be somewhat malleable, regulated by immune and metabolic reprogramming (22). Mutations in genes that regulate genomic integrity, such as *TP53,* can alter the intrinsic transcriptional program of tumor cells; however, genomic instability in the tumor can create even more dramatic changes by modulating transcriptional programs of heterotypic stromal cells in the tumor tissue, as well as in the composition of the niche (23,24).

Chromosome translocations and the resulting fusion genes are important contributors to the pathogenesis of cancer, particularly in sarcomas and hematopoietic malignancies (25). However, the nature and frequency of these events in canine HSA and human AS remains unclear. Here, we used next generation RNA sequencing (RNA-Seq) data to identify fusion genes in thirteen human ASs and 76 visceral HSAs originating from 74 dogs, and we investigated the relationship of these fusions to the mutational landscape of the tumor. We identified ten novel protein-coding fusion genes including *TEX2*-*PECAM1* and *ATP8A2*-*FLT1* in seven of thirteen human ASs, and two of the fusion-detected tumors showed mutations of *TP53* (R248Q and P250L). In canine HSAs, we found novel protein-coding fusion genes in a subset of the tumors of the angiogenic subtype. These fusion genes co-occurred with *TP53* mutations and were associated with gene enrichment for activated angiogenic pathways in the tumors. Our data suggest that genomic instability induced by mutations of *TP53* creates a permissive environment for fusion genes, with selection for angiogenic molecular programs in malignant vasoformative tumors. Our data also demonstrate that human AS and canine HSA maintain molecular programs that activate convergent signaling pathways to establish angiogenic phenotypes despite their genomic complexity.

## Materials and Methods

### Human tissue samples

Snap frozen and formalin fixed paraffin embedded (FFPE) tissues for human biospecimens were obtained from the University of Minnesota Biological Materials Procurement Network (UMN BioNet) and from the Cooperative Human Tissue Network (CHTN) under their standardized patient consent protocols. The demographic characteristics of human patients from whom we obtained ASs (*n* = 13) and normal tissue samples (*n* = 6) are summarized in **Supplementary Table S1**.

### Dog tissue samples

Seventy-six snap frozen and FFPE tissue samples were obtained from 74 dogs with HSAs. Frozen and FFPE tissues samples from 10 dogs with splenic hematomas, which are benign lesions with enlarged vascular spaces lined by endothelial cells, were used as controls. Samples were obtained as part of medically necessary diagnostic procedures and were used for research with owner consent. The origin of these samples was reported previously (14,21,26-28), or they were collected from dogs with HSA or with splenic hematomas at the Veterinary Medical Center, University of Minnesota. Procedures involving animal use were done with approval and under the supervision of the University of Minnesota Animal Care and Use Committee (protocols 1110A06186, 1507-32804A, 0802A27363, 1101A94713, 1312-31131A, and 1702- 34548A). The demographic characteristics of dogs (*n* = 74) from whom we acquired HSA and non-malignant splenic hematomas (*n* = 10) are summarized in **Supplementary Table S2**.

### Histological assessment

FFPE sections (4 μm) were stained with hematoxylin and eosin (H&E) and examined by veterinary pathologists to assign a histological diagnosis of canine HSA. Solid, capillary, cavernous, or mixed histological subtypes were assigned using accepted criteria (29); mitotic index (MI) was calculated per 1,000 cells in 5-10 random fields under 400X magnification (30). H&E slides were further reviewed for tumor content by two board-certified medical pathologists (ML and PM) (31), with the percent of sample containing viable nucleated cells corresponding to tumor recorded in a range of 0 to > 90% based on the planar surface of the sections. Diagnostic and histopathology reports of human tissues were provided by the specimen providers, the UMN BioNet and the CHTN.

### RNA isolation and generation of RNA-Seq libraries

Total RNA was isolated from tissue samples using the TriPure Isolation Reagent (Roche Applied Science, Indianapolis, IN, USA). The RNeasy Mini Kit (Qiagen, Valencia, CA, USA) was used for clean-up according to the manufacturer’s instructions. RNA-Seq from 74 canine HSA tissues is published (17,21,32,33) and an additional data set was generated from two canine HSA tissues and from 10 non-malignant splenic hematoma tissues. Total RNA was also extracted from 13 human AS tissues and from six normal tissues. Two μg of total RNA from each sample were quantified and assessed for quality; RNA-Seq libraries were generated as described (21) using the TruSeq RNA sample preparation kit (Illumina Inc., San Diego, CA). Sequencing was performed using HiSeq 2000 or 2500 systems (Illumina Inc.). Each sample was sequenced to a targeted depth of 20 - 80 million paired-end reads with mate-pair distance of 50 bp. Primary analysis and demultiplexing were performed using CASAVA software version 1.8.2 (Illumina Inc.) to verify the quality of the sequence data. The end result of the CASAVA workflow was demultiplexed into FASTQ files for analysis. Bioanalyzer quality control and RNA-Seq were performed at the University of Minnesota Genomics Center (UMGC) or at the Broad Institute.

### Bioinformatics analysis

The original FASTQ files prepared from thirteen human ASs and six non-malignant tissues were mapped to the human reference genome (GRCh38). The FASTQ files generated from 76 canine HSAs and ten non-malignant splenic hematomas were mapped to the dog reference genome (Canfam3.1). Sequencing quality was assessed by FastQC. The deFuse algorithm (34) was used to identify putative fusion events. To discriminate true fusion candidates from artifacts, we included fusion events with exon boundaries in both fusion partners and excluded events created from adjacent genes that showed breakpoint homology (>1). We also filtered highly recurrent fusion events that were found at implausible frequencies across tumor and non-malignant tissue samples (35) and transcription-induced chimeras. The split sequences of the fusion genes were validated by *de novo* assembly using Trinity (36). TranscriptsToOrfs and deFuse-Trinity tools verified the deFuse fusion predictions with Trinity-assembled transcripts and open reading frames. TopHat2 was used to generate BAM files, and the Integrative Genomics Viewer (IGV 2.3; Broad Institute, Cambridge, MA) was used to visualize the mate pair sequences of fusion genes. A protein translation tool in Expert Protein Analysis System (ExPASy; SIB Swiss Institute of Bioinformatics, Lausanne, Switzerland) was used to determine in-frame fusion proteins. Tumor purity and microenvironment scores were assessed using the bioinformatics tools ESTIMATE (37) and *xCell* (38).

### Reverse transcription polymerase chain reaction (RT-PCR) and Sanger sequencing

RT-PCR was performed to validate fusion transcripts identified by deFuse (39). Briefly, cDNA was synthesized using SuperScript^®^ VILO cDNA Synthesis Kit and Master Mix (Invitrogen). PCR amplification was performed using a conventional thermocycler with HotStarTaq DNA polymerase (Qiagen) or using a LightCycler® 96 (Roche Applied Science, Indianapolis, IN, USA) with FastStart SYBR Green Master Mix (Roche Applied Science) for quantitative real-time RT-PCR (40). PCR primer pairs used for fusion gene amplification are presented in the Results section. *GAPDH* was used as a control for RNA integrity and for the RT-PCR reactions. The forward and reverse primer sequences for *GAPDH* were 5’-GGA GTC CAC TGG CGT CTT CAC-3’ and 5’-GAG GCA TTG CTG ATG ATC TTG AGG-3’, respectively. Relative mRNA values were expressed as delta-Ct values normalized to *GAPDH*. Sanger sequencing was performed at the UMGC.

### Fluorescence *in situ* hybridization (FISH)

FISH was performed to detect *MYO16-PTK2* and *GABRA3-FLT1* fusion genes by designing FISH probes derived from the genome-anchored canine CHORI-82 bacterial artificial chromosome (BAC) library (41). Single locus probes were used for proximal *MYO16* at dog chromosome (CFA) 22:57,565,917-57,750,789 (clone 183H20), distal *MYO16* at CFA 22:57,750,801-57,967,880 (clone 385H13), and *PTK2* at CFA 13:35,302,679-35,483,060 (clone 451H13) with distinct fluorescent tags. For *GABRA3-FLT1* fusion, break-apart FISH probes were used for proximal *FLT1* at CFA 25:11,057,892-11,263,935 (clone 363B20) and distal *FLT1* at CFA 25:11,274,078-11,471,538 (clone 235H9). The PureLink® HiPure Plasmid Maxiprep Kit (Invitrogen) was used for BAC DNA extraction. For preparation and hybridization of FISH probes, BAC DNA probes were labeled by Nick Translation Kit (Abbott Molecular) using Green-500 dUTP, Orange-552 dUTP and Aqua-431 dUTP (Enzo Life Science). Labeled DNA was precipitated in COT-1 DNA, salmon sperm DNA, sodium acetate and 95% ethanol, then dried and resuspended in 50% formamide hybridization buffer. The Red-proximal *MYO16,* Green-distal *MYO16* and Aqua-*PTK2* probes were combined into one 3-color FISH probe for *MYO16-PTK2* fusion. The Red-proximal *FLT1* and Green-distal *FLT1* break-apart probes were applied for the split *FLT1* gene.

FFPE sections (4 μm) were processed according to the Dako IQFISH protocol; probes were applied to the slide and hybridized for 24 hours at 37°C in a humidified chamber. After hybridization, slides were washed and counterstained with DAPI. Fluorescent signals were visualized on an Olympus BX61 microscope workstation (Applied Spectral Imaging, Vista, CA) with DAPI, FITC, Texas Red and Aqua filter sets. FISH images were captured using an interferometer-based CCD cooled camera (ASI) and FISHView ASI software. A total of 200 interphase cells were examined for each sample. Non-malignant canine spleen tissues were used as controls for the FISH experiment.

### Validation of somatic mutations using RNA-Seq data

A pipeline was developed to identify the bases present at locations defined as somatic mutations in the Tumor-Normal exome calls (17). Briefly, RNA-Seq data were mapped using the STAR-Mapper (42) with STAR-FUSION mapping settings (43) to the Canfam3.1 or GRCh38 genome. BAM files generated by STAR were sorted and indexed using Samtools (44). Starting from a file containing somatic mutation locations and a file containing a list of BAM file locations, the pipeline uses Samtools (44) functions to identify the bases present at each location at each file and then reports whether a variant is found at that location. For inclusion within the variant file, at least one sample must have at least 3 reads that support the variant and that represent greater than 10% of all reads at that location.

### Gene expression profiling

CLC Bio Genomics Workbench 10 (CLC Bio, Aarhus, Denmark) and DESeq2 R packages were used to quantify gene expression and differential gene expression analysis as described (45). Briefly, paired-end RNA-Seq data with mate-pair distance of 50 bp in FASTQ format were imported, and sequencing quality was determined. Transcriptomics analysis was then performed to generate the expression level of each gene presented as total reads by mapping the sequencing reads to Canfam3.1 or GRCh38. Heatmaps and hierarchical clustering based on average linkage were visualized using Cluster 3.0, Morpheus (https://clue.io/morpheus), or R packages. GO Enrichment Analysis (46-48) or Ingenuity® Pathway Analysis software version 8.6 (Qiagen, Redwood City, CA) were used to define biological functions, canonical pathways, and upstream regulators associated with differently expressed genes (DEGs) between groups using Benjamini-Hochberg multiple testing corrections to evaluate significance. For gene expression profiling, unsupervised PCA and hierarchical clustering were performed to define subtypes of canine HSAs as described previously (21). Gene expression data of human sarcomas in The Cancer Genome Atlas (TCGA) database were also compared with our data sets.

### TMA generation and immunohistochemistry (IHC)

Canine TMA blocks were generated from 45 HSA tissues, including 32 tumors used for RNA-Seq and eight non-tumor tissues (six splenic hematomas and two non-malignant liver samples). Tissue cores of 1-mm diameter in quadruplicate from each sample were assembled in random order in four TMA blocks. One TMA block with mouse tissues was generated for staining controls. Immunostaining with CD31, Vimentin and Pan-Cytokeratin antibodies was evaluated to support tumor content estimates. A human TMA block was generated from ten AS tissues and six non-malignant tissues (submandibular gland, skin, breast adipose tissue, thigh skeletal muscle, spleen, and lung).

Unstained TMA sections (4 μm) were de-paraffinized and rehydrated using standard methods for IHC. All of the immunohistochemical assays, including validation for antibodies, were performed and optimized at the UMN BioNet Histology Laboratory or the Veterinary Diagnostic Laboratory at the University of Minnesota. Antibodies used for IHC are summarized in **Supplementary Table S3**. The immunostaining score assigned to each case was a semiquantitative assessment derived from the product of two integers, ranging from 0 to 3 and from 1 to 3, that respectively reflect the percentage of positive cells in a sample and the intensity of staining at high power magnification (400X) as described previously with some modifications (49). The percentage of positive cells was scored from 0 to 3+, where 0 reflected specific staining in <1% of the cells, 1+ reflected specific staining in >1% and <25% of the cells, 2+ reflected specific staining in 25-75% of the cells, and 3+ reflected specific staining in >75% of the cells. The intensity was assessed as weak (intensity score 1), moderate (intensity score 2), or strong (intensity score 3). Immunostaining results were scored (ranging from 0 to 9) by multiplying the percentage of positive cells (score 0-3) by the intensity (score 1-3).

### Statistical analysis

Chi-square or Fisher’s exact test, was performed for contingency tables analysis. Continuous values were analyzed by Welch’s (Heteroscedastic) T-test or Mann-Whitney U test. The statistical tests were two-tailed. Statistical analysis was performed using GraphPad Prism 6 (GraphPad Software, Inc., San Diego, CA). *P*-values are reported without inference of significance, consistent with the American Statistical Association’s Statement on Statistical Significance and P-Values (50).

## Results

### Novel protein-coding fusion genes are identified in human ASs and canine HSAs

Putative fusion gene events were identified from RNA-Seq data as paired-end sequence reads that mapped connecting two distant genes (**Supplementary Fig. S1A**). We identified novel in-frame protein-coding fusion transcripts for ten fusion events in 7 of 13 (53.8%) human ASs (**Fig. 1A**; **Table 1**). Two of the fusions were inter-chromosomal events and eight were intra-chromosomal events. The fusions included *TEX2-PECAM1,* which contained the gene that encodes CD31, and *ATP8A2-FLT1,* a kinase fusion gene that encodes the vascular endothelial growth factor receptor 1 (VEGFR1). None of the ten fusion events was seen in more than one tumor, and three of the seven fusion-positive tumors contained two distinct fusion events each.

**Figure 1.**
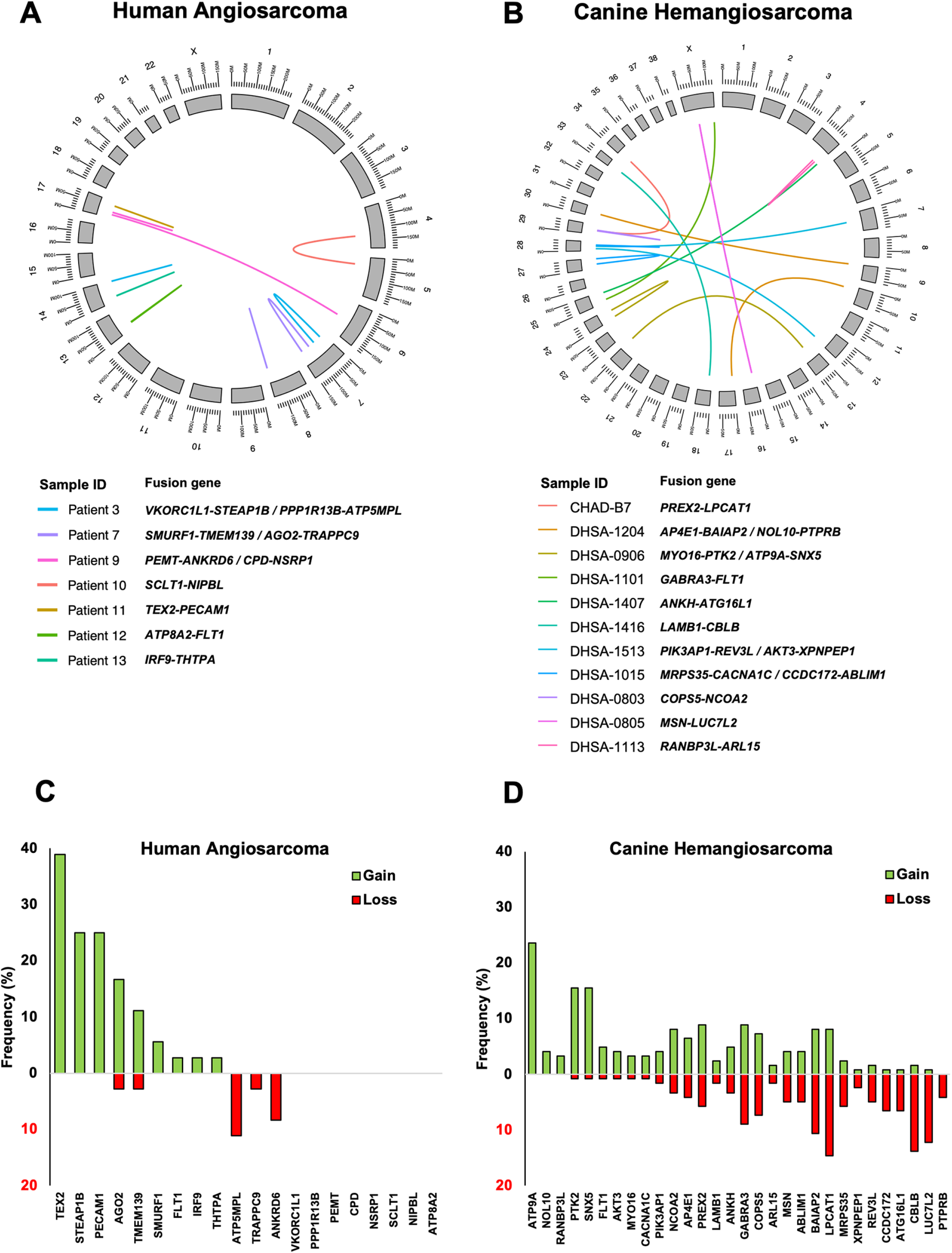
Identification of novel putative protein-coding fusion transcripts in human AS and canine HSA. The *Circos* plots visualize fusion genes identified in human ASs (**A**, *n* = 13) and canine HSA (**B**, *n* = 76). Bar graphs show DNA copy number alterations for each fusion partner gene using publicly available Exome-sequencing data in an independent dataset of human AS (**C**, *n* = 36; data retrieved from Ref (12)) and using oaCGH in a larger HSA dataset (**D**, *n* = 123; data retrieved from Ref (19)).

**Table 1.** Putative fusion genes identified in transcriptomic data of human ASs

| Patient ID | Gene 1 | Gene 2 | Putative fusion gene | Gene 1 chromosome | Gene 2 chromosome | Gene 1 fusion location | Gene 2 fusion location | Fusion Type | Gene 1 Ensembl ID | Gene 2 Ensembl ID | Genomic break position in gene 1 | Genomic break position in gene 2 | Somatic variants |
| --- | --- | --- | --- | --- | --- | --- | --- | --- | --- | --- | --- | --- | --- |
| Patient 1 | - | - | - | - | - | - | - | - | - | - | - | - | - |
| Patient 2 | - | - | - | - | - | - | - | - | - | - | - | - | - |
| Patient 3 | VKORC1L1 | STEAP1B | VKORC1L1-STEAP1B | 7 | 7 | coding | coding | Intra-chromosomal | ENSG00000196715 | ENSG00000105889 | 65873565 | 22419836 | - |
|  | PPP1R13B | ATP5MPL | PPP1R13B-ATP5MPL | 14 | 14 | coding | coding | Intra-chromosomal | ENSG00000088808 | ENSG00000156411 | 103847299 | 103915189 | - |
| Patient 4 | - | - | - | - | - | - | - | - | - | - | - | - | - |
| Patient 5 | - | - | - | - | - | - | - | - | - | - | - | - | HRAS |
| Patient 6 | - | - | - | - | - | - | - | - | - | - | - | - | - |
| Patient 7 | SMURF1 | TMEM139 | SMURF1-TMEM139 | 7 | 7 | coding | intron | Intra-chromosomal | ENSG00000198742 | ENSG00000178826 | 99143726 | 143286424 | TP53 |
|  | AGO2 | TRAPPC9 | AGO2-TRAPPC9 | 8 | 8 | coding | coding | Intra-chromosomal | ENSG00000123908 | ENSG00000167632 | 140635485 | 140451383 |  |
| Patient 8 | - | - | - | - | - | - | - | - | - | - | - | - | NRAS |
| Patient 9 | PEMT | ANKRD6 | PEMT-ANKRD6 | 17 | 6 | coding | utr5p | Inter-chromosomal | ENSG00000133027 | ENSG00000135299 | 17577027 | 89433375 | - |
|  | CPD | NSRP1 | CPD-NSRP1 | 17 | 17 | coding | coding | Intra-chromosomal | ENSG00000108582 | ENSG00000126653 | 30461311 | 30172542 | - |
| Patient 10 | SCLT1 | NIPBL | SCLT1-NIPBL | 4 | 5 | coding | coding | Inter-chromosomal | ENSG00000151466 | ENSG00000164190 | 128888679 | 37057186 | - |
| Patient 11 | TEX2 | PECAM1 | TEX2-PECAM1 | 17 | 17 | utr5p | utr5p | Intra-chromosomal | ENSG00000136478 | ENSG000000261371 | 64263168 | 64390763 | - |
| Patient 12 | ATP8A2 | FLT1 | ATP8A2-FLT1 | 13 | 13 | coding | coding | Intra-chromosomal | ENSG00000132932 | ENSG00000102755 | 25968679 | 28357685 | - |
| Patient 13 | IRF9 | THTPA | IRF9-THTPA | 14 | 14 | coding | coding | Intra-chromosomal | ENSG000000213928 | ENSG000000259431 | 24164134 | 23558695 | TP53 |

In canine HSA, we found fifteen novel protein-coding fusion genes in eleven of 76 tumors (14.5%) (**Fig. 1B**; **Table 2**). Ten of the fusions were inter-chromosomal events and five were intra-chromosomal events. None of the fifteen fusion events was seen in more than one tumor, and four of the eleven fusion-positive tumors involved two distinct fusion genes each. One fusion partner in four of the translocations encoded either a protein kinase or a protein phosphatase associated with angiogenic signaling (*MYO16-PTK2, GABRA3-FLT1, AKT3-XPNPEP1,* and *PTPRB-NOL10).* Eight of the fusion genes were associated with kinase signaling or kinase binding activity, such as PI3 kinase signaling, a MAP kinase, a receptor tyrosine kinase, a protein serine/threonine kinase, and an NAD+ kinase. Gene ontology annotations of the fusion partners for every translocation are described in **Supplementary Table S4** for human AS and **Supplementary Table S5** for canine HSA. Fusion genes were not present in any of the human non-malignant tissues (*n* = 6) or canine hematomas (*n* = 10) examined. Conserved driver translocations such as *BCR-ABL* and *MYC-IGH* that are present in both human and canine chronic myelogenous leukemia and Burkitt lymphoma, respectively (51), were not identified in human AS and canine HSA. All the fusion events identified in the seven human ASs and eleven canine HSAs involved different gene pairs, with the exception of the *FLT1* gene, which created fusions with a different partner gene in one case from each species.

**Table 2.**
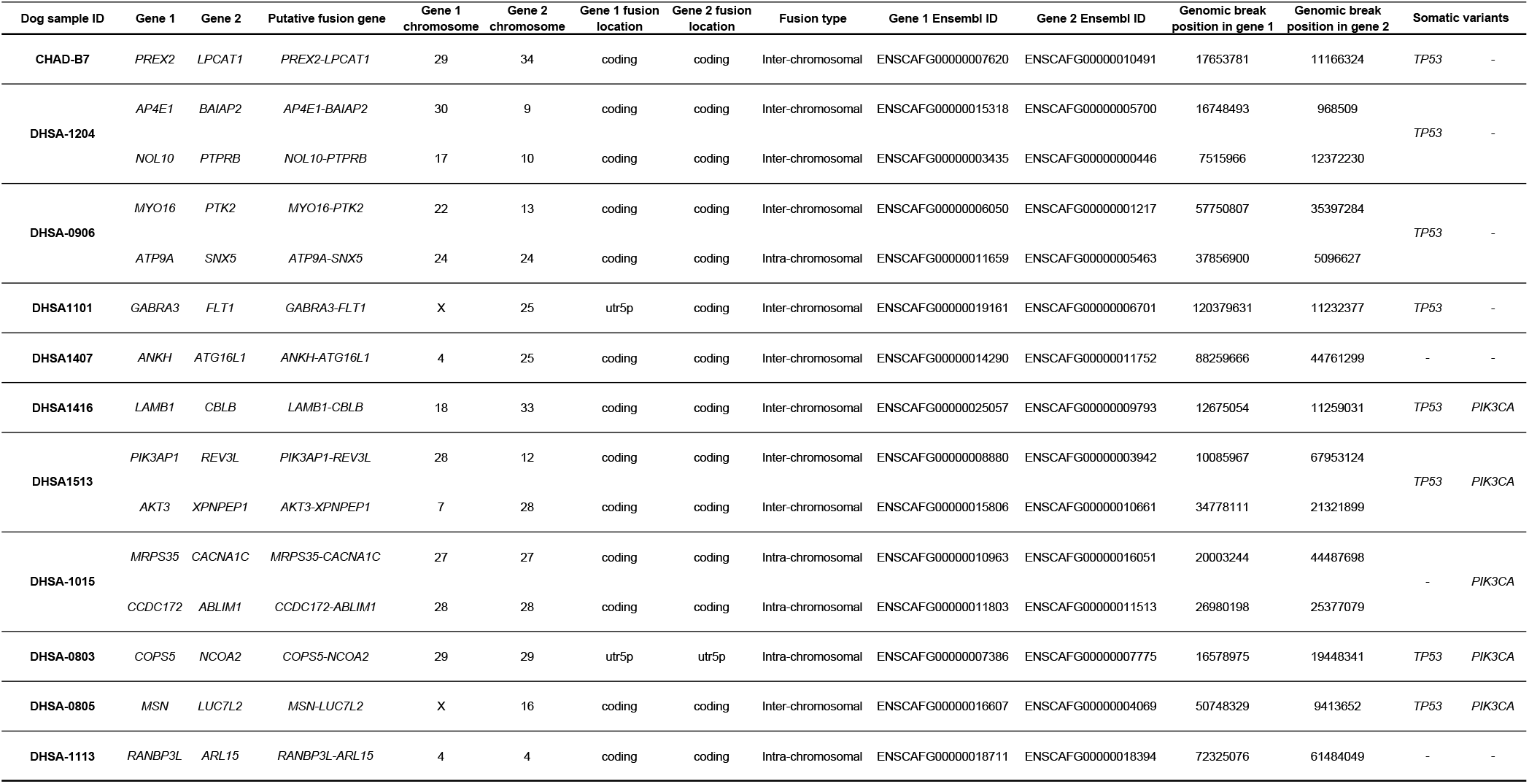
Putative fusion genes identified in transcriptomic data of canine HSAs

To determine whether the non-tumor components in the tumor tissue affected the detection of fusion genes, we quantified tumor content histologically and bioinformatically in canine HSAs **(Supplementary Fig. S2A - E**). Seventy of the 76 HSA samples were histologically evaluated, and the tumor content was not different between HSA samples with fusion events (*n* = 11) and those without fusion events (*n* = 59). We used two independent bioinformatic tools, *xCell* and ESTIMATE, to predict stromal and immune cell components, and these algorithms generated consistent output scores (Pearson’s *R* = 0.84; *R*^2^ = 0.71; *P* < 0.00001) that showed the presence of fusion genes was not associated with tumor purity. The detection of fusion genes was also independent of sequencing depth (**Supplementary Fig. S2F**). To rule out artifacts from the computational process, we validated the presence of the inter-chromosomal fusion gene, *SCLT1-NIPBL,* in the original human AS sample where it was identified, in two additional samples where it was undetectable based on sequencing data, and in a non-malignant tissue sample. The fusion transcript was detectable by quantitative real time RT-PCR amplification; PCR primer pairs were designed to amplify putative split sequences (up to 200 base pairs) involving the breakpoints identified by deFuse (**Supplementary Fig. S3A**). We confirmed that the junction sequences between the two genes producing the new fusion event were amplified by PCR (**Supplementary Fig. S3B**). Four representative fusion transcripts found in canine HSAs (*MYO16-PTK2; AKT3-XPNPEP1; AP4E1-BAIAP2; NOL10-PTPRB)* were also detected, but only in the respective cases where they were identified in the sequencing data (**Supplementary Fig. S3C** - **E**). Each PCR amplification product was verified by Sanger sequencing. We then used RT-PCR to evaluate RNA-Seq data from 63 canine tissue samples (53 HSAs and 10 hematomas) for the presence of these four fusion transcripts. The results were consistent between RNA-Seq and PCR, as we found neither false-positive nor false-negative events in the samples tested (**Supplementary Table S6**).

### Fusion genes are associated with DNA copy number variations

We then determined if any of the fusion partner genes identified in our analysis were associated with DNA copy number alterations. Publicly available whole Exome-sequencing data generated from an independent data set of 36 human patients with ASs was used (12). Copy number variations were found in twelve of the twenty (60%) fusion partner genes: nine genes were amplified, and five genes were deleted (**Fig. 1C**; **Supplementary Fig. S4)**. *TEX2* (39%), *STEAP1B* (25%) and *PECAM1* (25%) were the top three genes where copy number gains occurred most frequently. For canine HSA, we used oligonucleotide array comparative genomic hybridization (oaCGH) in a larger HSA dataset (*n* = 123) (19). Copy number gains were observed in 29 of the 30 (96.7%) fusion partners, and copy number losses were observed in 27 of the 30 (90.0%) fusion partners. Protein kinase-encoding genes, *PTK2, FLT1,* and *AKT3* revealed a higher frequency of copy number gain: 15.5% gain vs 0.8% loss for *PTK2;* 4.9% gain vs 0.8% loss for *FLT1;* and 4.1% gain vs 0.8% loss for *AKT3,* suggesting that copy number alterations leading to dysregulation of downstream kinase signaling contribute at least partly to the angiogenic program in a subset of canine HSAs (**Fig. 1D**).

### Chromosomal translocations resulting in fusion genes are detectable in canine HSAs

We performed FISH to confirm that fusion genes were generated by chromosome translocations. We chose representative inter-chromosomal fusions, *MYO16-PTK2* and *GABRA3-FLT1* for cytogenetic validation because PTK2 and VEGFR1 are key molecules that regulate pathogenic signaling in vascular cancers, including canine HSA (52). **Fig. 2A** illustrates the predicted structure of *MYO16-PTK2* inter-chromosomal fusion between CFA 22 and CFA 13, based on deFuse and Sanger sequencing data (**Fig. 2B**). The predicted fusion gene comprises exons 1-32 of *MYO16* (CFA 22) and exons 12-31 of *PTK2* (CFA 13), with the putative junction joining *MYO16* exon 32 and *PTK2* exon 12. Breakage occurs between exons 32 and 33 of *MYO16* at CFA 22:57,750,807 bp, and between exons 11 and 12 of *PTK2* at CFA 13:35,397,284 bp. Independent FISH probes identifying the association between proximal and distal *MYO16* and the breakpoint of *PTK2* (**Fig. 2C**) confirmed the presence of the *MYO16-PTK2* fusion gene between CFA 22 and 13 in archival FFPE samples from the same dog tumor. The *MYO16-PTK2* fusion was identified by deFuse and RT-PCR (**Fig. 2D**). The t(CFA 13;CFA 22) translocation was present in interphase nuclei of 16.8% of the tumor cells, with a smaller subpopulation showing amplification of the fusion (**Fig. 2E**).

**Figure 2.**
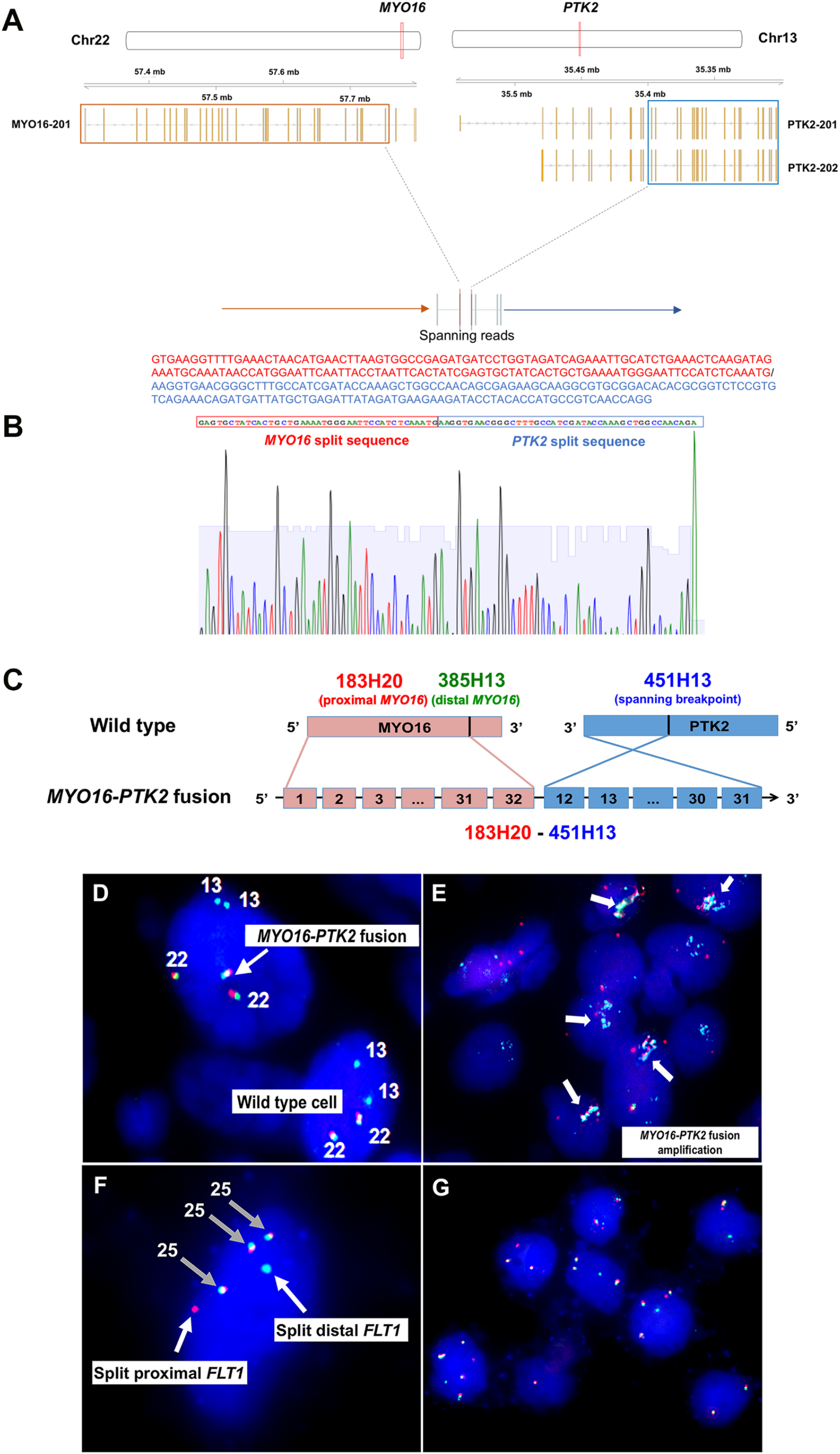
Validation of fusion genes in canine HSA. **A,** *MYO16-PTK2* fusion gene track and visualization of the breakpoint in UCSC Genome Browser (Canfam3.1). **B,** Sanger sequencing result of the PCR product for *MYO16-PTK2* fusion gene. **C,** Schematic illustration of the putative *MYO16-PTK2* fusion gene and designed FISH probes. **D,** Detection of the *MYO16- PTK2* fusion gene on primary canine HSA tissue by FISH. In wild type cells an association between BAC clones 183H20 (red) and 385H13 (green) can be appreciated, both showing independent localization from 451H13 (aqua). A portion of tumor cells shows a breakage within 451H13, with one half of that signal associating with 183H20, and independent of the localization of 385H13 indicating the existence of the *MYO16-PTK2* fusion at the genomic level. **E,** Arrows indicate the amplification of the *MYO16-PTK2* fusion gene. **F,** Detection of the *GABRA3-FLT1* fusion gene by FISH. The *GABRA3-FLT1* fusion is identified by break-apart FISH probes for proximal *FLT1* (clone 363B20; red) at CFA 25 and distal *FLT1* at CFA 25 (clone 235H9; green). Split *FLT1* genes indicate the fusion event identified by single color signal (white arrows). Dual colors represent the intact *FLT1* gene (grey arrows). **G,** DNA amplification of *FLT1* gene.

We used break-apart FISH to validate the presence of the *GABRA3-FLT1* fusion gene (**Fig. 2F**). Split *FLT1* probes were found in 36.7% of tumor cells in archival FFPE samples from the dog tumor in which the *GABRA3-FLT1* fusion was identified by deFuse and RT-PCR. Interestingly, in this tumor the intact *FLT1* gene showed consistent amplification (up to four copies), suggesting Flt-1 (also known as VEGFR1) activation in this tumor might have occurred through multiple mechanisms (**Fig. 2G**). We next used FISH analysis to assess recurrence of the *MYO16-PTK2* fusion in a tissue microarray (TMA) comprised of 45 visceral HSAs and eight non-malignant tissues (six spleens; two livers). The *MYO16-PTK2* fusion was once again present in the sample from the canine tumor in which it was discovered, but it was not seen in any other sample on the TMA. We also performed FISH to detect the *ATP8A2-FLT1* fusion in human AS using a break-apart *FLT1* probe, but the fusion was undetectable in our FFPE sample. This might have been due to the small number of tumor cells that were likely to contain the fusion event in a heterogeneous clonal population. Since none of the fusion transcripts identified in our cohorts of human and canine tumors were recurrent, we sought to determine if the fusion events were associated with other genetic and molecular programs.

### Fusion genes and somatic variants in human ASs enrich angiogenic gene signatures

To examine genomic aberrations associated with the fusion genes, we determined somatic variations and gene expression profiles using RNA-Seq data. In human ASs, *TP53* mutations (R248Q and P250L) were observed in two of thirteen human ASs, which also had fusion genes (*SMURF1-TMEM139* and *AGO2-TRAPPC9* in one tumor, *IRF9-THTPA* in the other tumor). *NRAS* (Q61L; *n* = 1) or *HRAS* (Q61L; *n* = 1) mutations were also detected, and both were present in tumors that did not have fusion genes or *TP53* mutations (**Fig. 3A**; **Table 1**). The RNA-Seq data did not provide evidence of mutations in *PIK3CA, PTEN,* or *KRAS* in this group of thirteen ASs.

**Figure 3.**
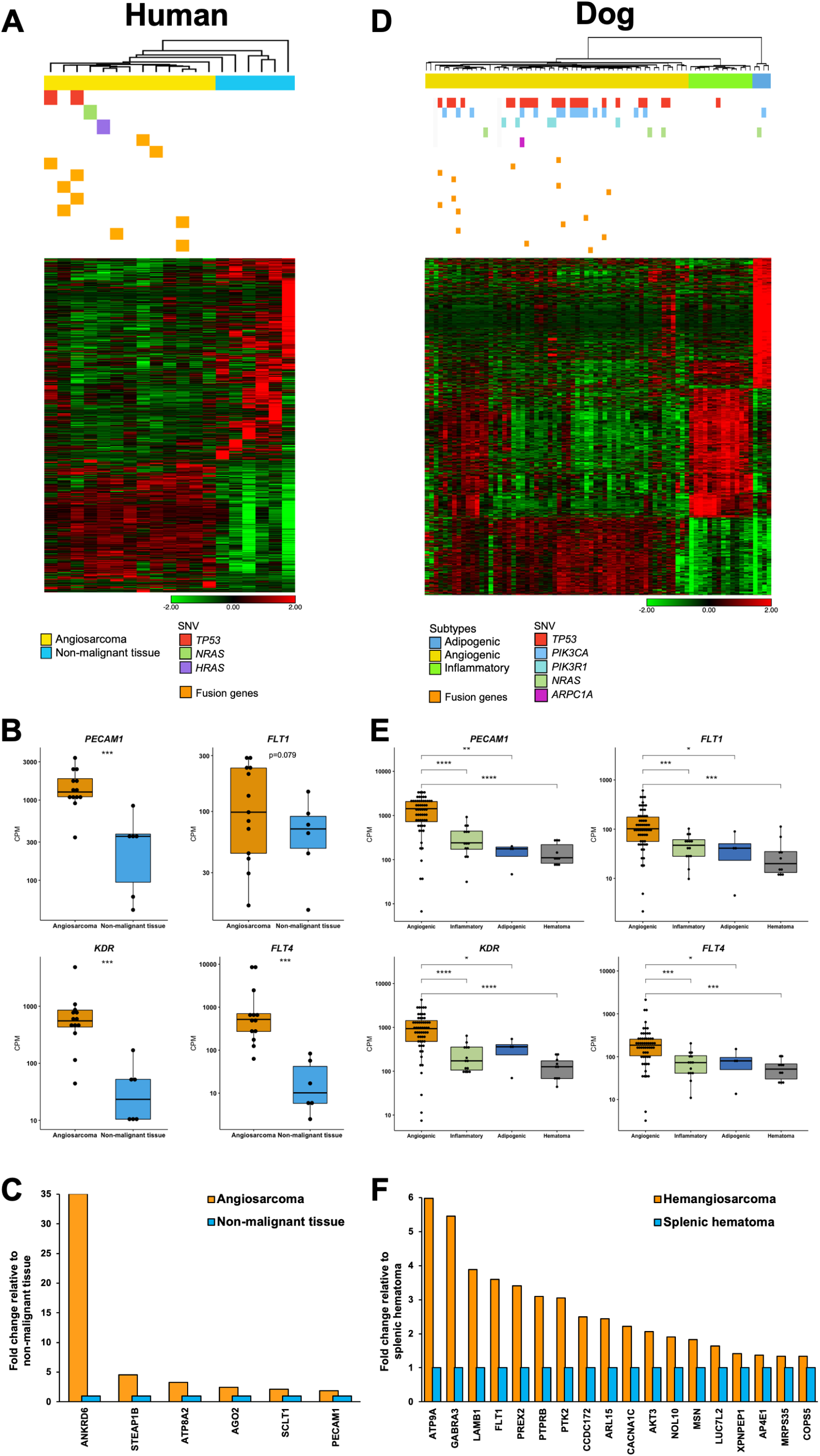
Fusion genes, mutations of somatic variants, and molecular subtypes of human AS (A-C) and canine HSA (D-F). **A,** 1,237 differentially expressed genes were identified between human ASs (*n* = 13) and non-malignant tissue samples (*n* = 6) (False Discovery Rate or FDR *P* < 0.05): 490 genes were upregulated and 747 genes were downregulated in ASs. Ten fusion genes are marked as yellow bars in seven AS samples. Somatic variations in *TP53 (n* = 2), *NRAS (n* = 1), and *HRAS (n* = 1) were found in four ASs. **B,** Box plots show gene expression of *PECAM1, FLT1, KDR,* and *FLT4* representing vasculogenic and angiogenic functions in human ASs and non-malignant tissues (two-tailed Mann-Whitney test). ***, *P* < 0.001. **C**, Bar graphs show the relative expression of genes in human ASs normalized to the expression of non-malignant tissues. Six of 20 genes where the *P*-value was less than 0.05 are displayed (two-tailed Welch’s T-test). **D,** Heatmap illustrates 1,477 significant differentially expressed genes among three subtypes of canine HSA (*n* = 76) (FDR *P* < 0.001; Fold change > 3). Somatic variant analysis identified mutations in *TP53 (n* = 24), *PIK3CA (n* = 16), *PIK3R1 (n* = 5), *NRAS (n* = 4), and *ARPC1A (n* = 1). Grey bars indicate unavailable somatic variants data. Fifteen fusion genes are marked as yellow bars in 11 HSA samples. **E,** Box plots display gene expression of *PECAM1, FLT1, KDR,* and *FLT4* representing vasculogenic and angiogenic functions in subtypes of HSA and non-malignant hematomas (two-tailed Mann-Whitney test). ****, *P* < 0.0001; ***, *P* < 0.001; **, *P* < 0.01; *, *P* < 0.05. **F**, Bar graphs show the relative expression of genes in canine HSA normalized to the expression of hematomas. Eighteen of 30 genes where the P-value is less than 0.05 are displayed (two-tailed Welch’s T-test). Heatmaps (**A** and **D**) show unsupervised hierarchical clustering (average linkage). Heatmap colors display mean-centered fold change expression following log2 transformation. Upregulated genes are presented in red and downregulated genes are shown in green.

We established transcriptomic profiles of human ASs to identify molecular traits that regulate global gene expression. We identified 1,237 differentially expressed genes between ASs (*n* = 13) and non-malignant controls (*N* = 6) (FDR *P*-value < 0.05): 490 genes were upregulated and 747 genes were downregulated in ASs. Biological functions and pathway analysis revealed that upregulated genes in ASs were associated with cancer, angiogenesis, vasculogenesis, and development of vasculature (*P* < 0.00001) (**Supplementary Table S7**). Additionally, we performed cell type enrichment analysis using the *xCell* tool (38) to predict relative populations of cellular components that comprised the AS tissues (**Supplementary Fig. S5**). We compared the cell type signature of AS with that of sarcomas (*n* = 263) from the TCGA database, which did not include AS. The results showed that gene signatures associated with endothelial cells and activated dendritic cells were highly enriched in ASs, while other sarcomas in the TCGA revealed gene enrichment of smooth muscle cells. Gene expression profiles of non-malignant samples indicated distinct tissue-specific patterns of submandibular gland, skin, breast adipose tissue, thigh skeletal muscle, spleen, and lung. The ASs showed upregulation of key angiogenic genes such as *PECAM1 (CD31), FLT1 (VEGFR1), KDR (VEGFR2),* and *FLT4 (VEGFR3)* compared to non-malignant tissues (**Fig. 3B**). In addition, 6 of 20 (30.0%) fusion partners (including *PECAM1)* showed a higher level of expression in the tumors compared to non-malignant tissues (*P* < 0.05; fold change in a range from 1.9 to 35.0) (**Fig. 3C**). Collectively, our data showed enriched angiogenic molecular programs were present in human ASs. Furthermore, both tumors with *TP53* mutations also harbored fusion genes.

### Fusion genes that co-occur with mutations of *TP53* are present exclusively in angiogenic canine HSAs

In canine HSAs, *TP53 (n* = 24/74; 32.4%), *PIK3CA (n* = 16/74; 21.6%), *PIK3R1 (n* = 5/74; 6.8%), and *NRAS (n* = 4/74; 5.4%) transcripts showed recurrent mutations that were consistent with those identified using tumor:normal Exome-sequencing (17). We found associations between mutations of *TP53* (*TP53*^mt^) and *PIK3CA (PIK3CA^mt^)* and fusion genes (Fusion^+^) (**Supplementary Table S8**). Specifically, *TP53*^mt^ commonly co-occurred with *PIK3CA*^mt^ (*P* = 0.004) and with fusion genes (*P* = 0.004); and Fusion^+^ tumors were seen in the tumors with *TP53*^mt^ or *PIK3CA*^mt^ (*P* = 0.005) more frequently than would be expected by random chance. When fusion genes co-occurred with *PIK3CA^mt^,* they invariably co-occurred with mutations of *TP53 (P* = 0.037), and they were not associated with *PIK3CA^mt^* alone (*P* = 0.324).

Next, we sought to determine if fusion genes were preferentially associated with specific molecular subtypes of canine HSA. We previously defined distinct angiogenic, inflammatory, and adipogenic molecular subtypes of canine HSA (21). To further validate this classification, we applied unsupervised principal component analysis (PCA) and hierarchical clustering to identify distinct groups in the sample cohort from this study (**Supplementary Fig. S6**). Our results show that the three molecular subtypes (58 angiogenic, 14 inflammatory, and 4 adipogenic) were reproducibly identified in the current dataset, as illustrated in the heatmap of 1,477 DEGs (FDR *P* < 0.001; fold change > |3|) shown in **Fig. 3D**. Interestingly, fusion genes were present only in tumors of the angiogenic HSA subtype (*P* = 0.046). Likewise, *TP53* mutations were identified in 23 of 56 (41.1%) angiogenic HSAs, and in one of 18 tumors from the two other molecular subtypes (*P* = 0.008). The somatic variants of *TP53, PIK3CA, PIK3R1,* and *NRAS* were found in 33 of 56 (58.9%) angiogenic HSAs, and in 3 of 18 (16.7%) tumors from the two other subtypes (*P* = 0.002). The angiogenic subtype of HSA also showed upregulation of *PECAM1 (CD31), FLT1 (VEGFR1), KDR (VEGFR2),* and *FLT4 (VEGFR3)* compared to the other two HSA subtypes and to non-malignant hematomas (**Fig. 3E**). We found that five of 32 dogs (16%) with angiogenic HSA lived longer than five months, while four of 10 dogs (40%) with inflammatory HSA survived longer than that (**Supplementary Fig. S7**). Eighteen of 30 (60.0%) fusion partners, including protein kinase-encoding genes such as *FLT1, PTK2,* and *AKT3,* showed higher levels of expression in HSAs compared to non-malignant controls (*P* < 0.05; fold change in a range from 1.3 to 6.0) (**Fig. 3F**). Neither breed, sex, neuter status, age, nor affected organs were associated with the presence of fusion genes (**Supplementary Fig. S8**). There was also no association between the fusion events and histological subtype or mitotic index (**Supplementary Table S9**).

Next, we analyzed DEGs (FDR *P* < 0.05; fold change > |2|) and gene pathways to examine gene signatures enriched in HSAs that had both fusion genes and *TP53* mutations. We classified tumors according to their mutations as summarized in **Supplementary Table S10**. Co-occurrence of fusion genes with *TP53* mutations (i.e., *TP53*^mt^/Fusion^+^/*PIK3CA*^wt^ or PF tumors) was associated with angiogenic and vascular signaling with enrichment of genes in pathways such as PI3K, VEGF, and PDGF (**Supplementary Table S11 - S13**). Thirteen genes that were commonly enriched in PF tumors were associated with activation of WNT3A as an upstream regulator (**Supplementary Fig. S9**). **Fig. 4** illustrates a model integrating the data from these findings to highlight potential pathogenetic contributions of fusion genes and recurrent mutations in canine HSA.

**Figure 4.**
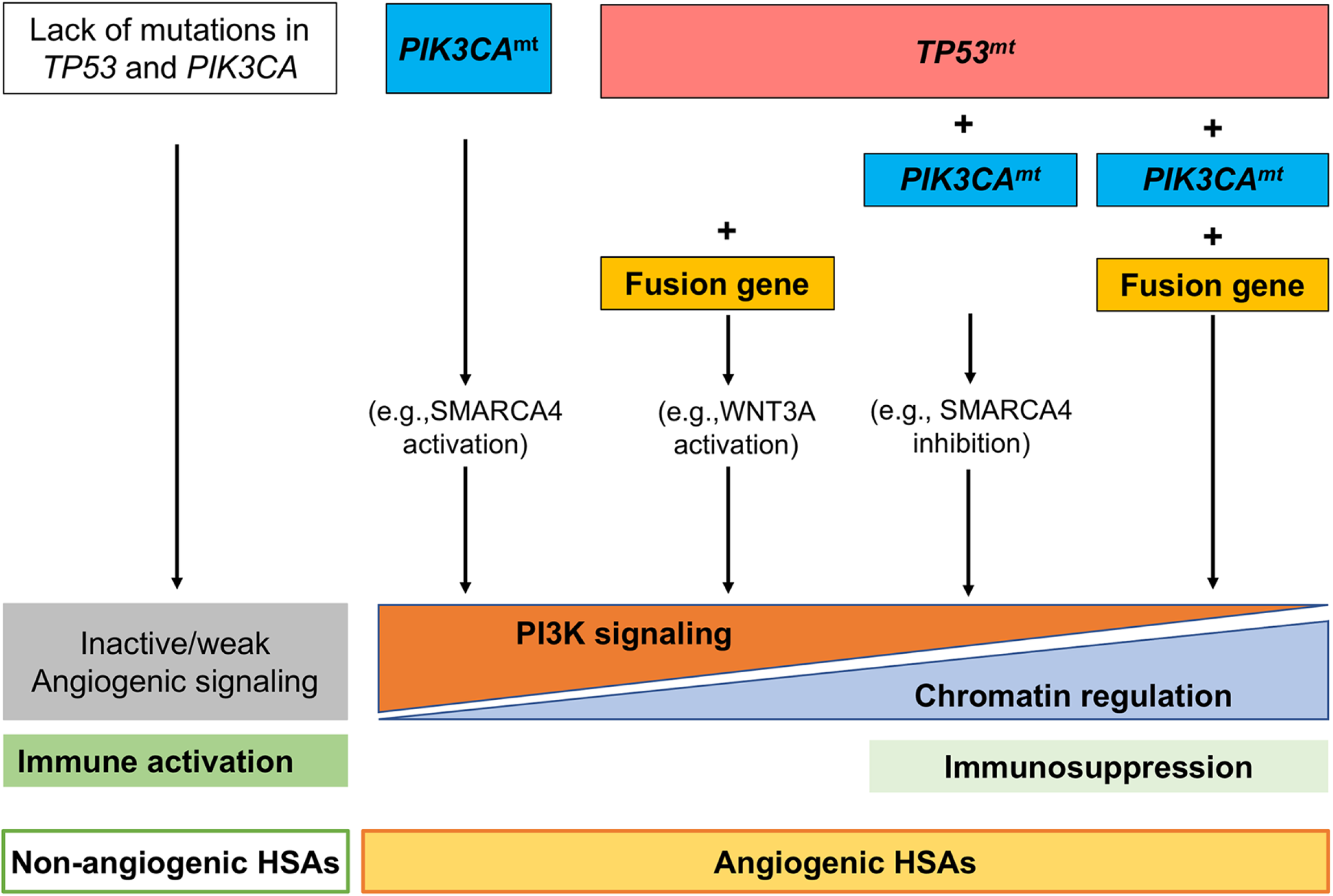
Hypothetical model for angiogenic pathogenesis of canine HSA.

### Human ASs and canine HSAs establish molecular programs that activate convergent signaling pathways

To determine if the genetic and molecular features of human AS and canine HSAs contributed to the activation of functional pathways, we performed IHC in eleven human AS tissues and in 44 canine HSAs (**Fig. 5**; **Supplementary Tables S14** and **S15**). First, we evaluated the effects of *TP53* mutation on the presence and location of p53, phospho-p53 (Ser15), and phospho-p53 (Ser20) (**Fig. 5A** and **B)**. In human ASs, nuclear expression of p53 was found in all of eleven (100%) tumors, showing various levels of expression. Nuclear expression of phospho-p53 (Ser15) was seen in seven of eleven (63.6%) tumors, showing low expression in five of seven (71%) tumors (IHC score ≤ 3). Nuclear and cytoplasmic expression of phospho-p53 (Ser20) protein was detected in all eleven (100%) tumors, and seven of those showed high expression (IHC score ≥ 7). In canine HSAs, p53 protein was localized to the nucleus in 34 of 44 (77%) tumors with various levels of expression. Immunoreactivity of phospho-p53 (Ser15) was observed in the nuclei of tumor cells in 38 of 40 (95%) cases, with 34 (90%) showing low or medium expression (IHC score ≤ 6). Nuclear and cytoplasmic expression of phospho-p53 (Ser20) was seen in all of 40 HSAs (100%), with 32 tumors (80%) showing high levels of expression. These data revealed that patterns of p53 and activated p53 were comparable in human ASs and canine HSAs; especially, p53 was strongly phosphorylated at residue Ser20 in tumors from both species. We found no association between phosphorylated p53 and *TP53* mutations or fusion genes (**Supplementary Fig. S10**), suggesting that DNA damage and cellular stress are widespread among these tumors, and they are likely to activate p53-mediated repair mechanisms independent of these genetic alterations.

**Figure 5.**
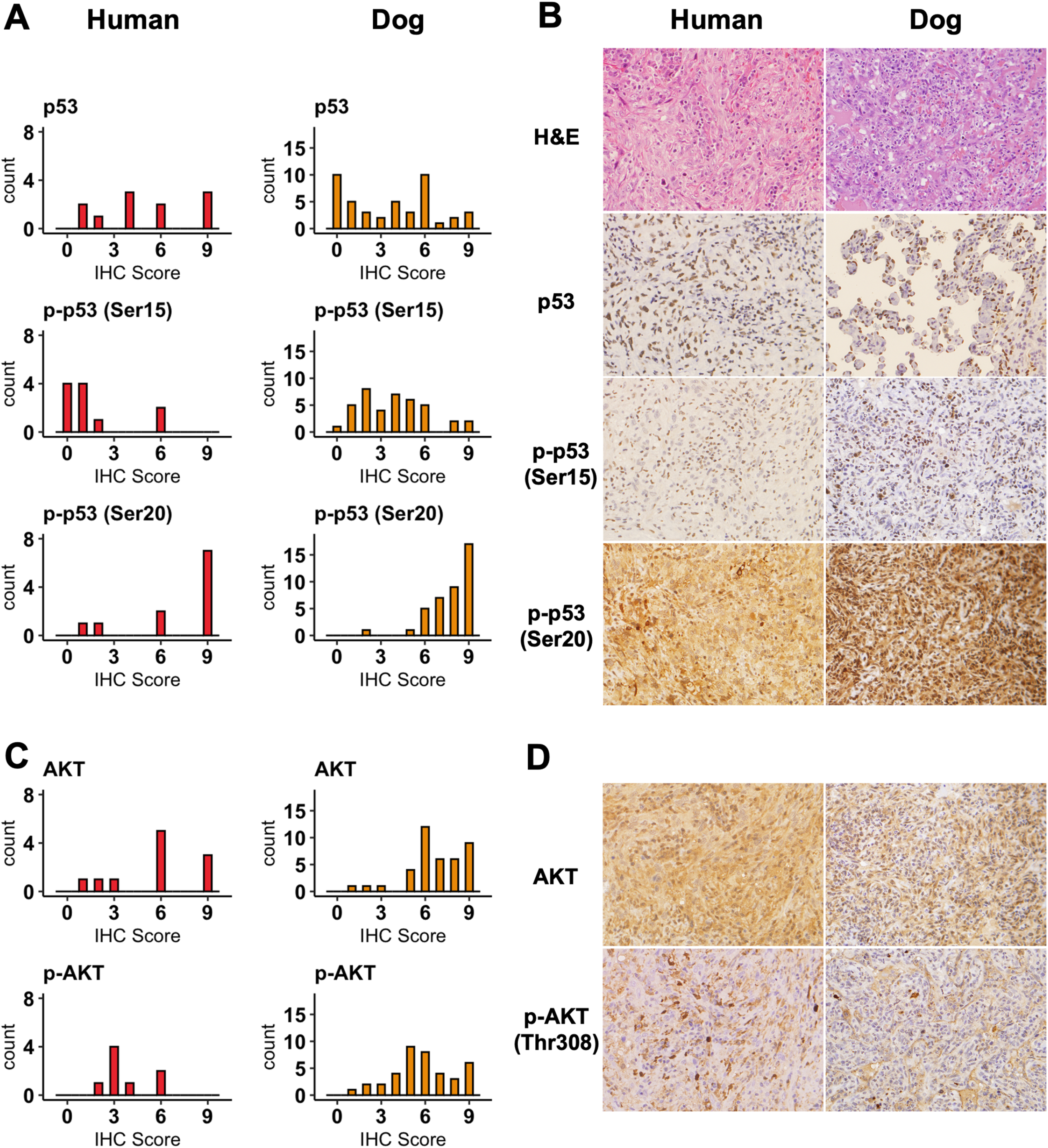
Immunohistochemical expression of p53 and AKT proteins in human AS and canine HSA. **A,** Bar graphs show the number of cases (y-axis) plotted as a function of IHC scores (x-axis) for staining with anti-p53, anti-phosphorylated (p)-p53 (Ser15), and anti-p-p53 (Ser20) antibodies in human ASs (Left panel) and canine HSAs (Right panel). **B,** Representative photomicrographs show IHC staining of p53, p-p53 (Ser15), and p-p53 (Ser20) in human AS (Left panel) and canine HSA tissues (Right panel). Bar graphs (**C**) for IHC scores of AKT and p-AKT (Thr308) proteins and representative photomicrographs (**D**) are also displayed for human ASs and canine HSAs. H&E = hematoxylin and eosin stain. IHC staining (Horseradish peroxidase and hematoxylin counterstain). 200X magnification.

The PI3K/AKT/mTOR signaling pathway is important for regulation of angiogenic, vascular, and energetic functions. To assess whether PI3K mutations resulted in higher levels of downstream pathway activation, we evaluated expression of AKT and phospho-AKT proteins in human ASs and canine HSAs (**Fig. 5C** and **D**). Expression of nuclear and cytoplasmic AKT protein was observed in all eleven human ASs (100%) with eight of eleven (73%) tumors showing medium or high expression. Expression of phospho-AKT (Thr308) was evaluated in eight tumors; all of them (100%) showed weak or medium levels of expression. Similarly, AKT protein was detectable in the nucleus and cytoplasm of all forty (100%) canine HSAs with 37 (93%) showing medium or high expression. Phospho-AKT (Thr308) was detected in the nuclei and cytoplasm of all evaluable 39 (100%) canine HSAs, with 34 (87%) expressing medium or high level of the protein. Strong AKT immunoreactivity was also seen in scattered stromal cells in both human AS and canine HSA. Neither mutations of *TP53, PIK3CA, PIK3R1* nor the presence of fusion genes were associated with expression of AKT and phospho-AKT in human or canine tumors (**Supplementary Fig. S11**). Since *PIK3CA* and *PIK3R1* mutations were undetectable in this set of eleven human ASs, we examined expression of mTOR and phospho-mTOR (Ser2448) proteins as surrogates to confirm activation of their downstream pathways in these tumors. mTOR protein was observed in the nuclei and cytoplasm of all eleven tumors (IHC score ≥ 6), and it was not associated with the presence of *TP53* mutations or fusions (**Supplementary Fig. S12A** and **B**). However, nuclear and cytoplasmic expression of phospho-mTOR was higher in ASs that had *TP53* mutations or that had fusion genes than it was in tumors without one of these genetic changes (**Supplementary Fig. S12C** and **D**). In summary, the immunostaining data suggest that human AS and canine HSA have comparable activation of the p53 and PI3K/AKT/mTOR pathways, and these events are largely independent of their mutational states. Our results further suggest that these vasoformative tumors from both species activate convergent signaling pathways that contribute to their final architecture and organization with predictable enrichment of angiogenic gene signatures.

## Discussion

For this study, our objective was to identify novel fusion genes in human ASs and spontaneous canine HSAs. We showed that novel protein-coding fusion genes were identified in approximately 50% of human ASs of which two had *TP53* mutations. In canine HSAs, protein-coding fusion genes were detectable in ~15% of tumors, and those were associated with p53 deficiency and enrichment of angiogenic gene signatures. Our data suggest that convergent molecular mechanisms associated with p53 inactivation and enhanced PI3K/AKT/mTOR signaling pathways are operational in genomically complex human ASs and canine HSAs.

In the past decade, advances in next-generation sequencing and bioinformatics have enabled genome-wide identification of unbiased cancer-associated fusion transcripts in a variety of tumor types. Previous studies have reported 7,887 fusion transcripts identified across thirteen tumor types in TCGA datasets and 9,928 fusion genes with a 3% recurrence rate in the Mitelman Database of Chromosome Aberrations and Gene Fusions in Cancer (53-55). These findings illustrate the complexity of the cancer-associated fusion gene landscape, showing a relatively high rate of fusions with low recurrence, possibly arising from catastrophic chromosome rearrangements by chromothripsis (56) and chromoplexy (57). Despite this relatively high frequency of fusion genes, a solution to define their pathogenic significance remains elusive. One key finding from this work was that protein-coding fusion genes co-occurred with mutations of *TP53* in the angiogenic molecular subtype of canine HSA, suggesting that genomic instability might create a predisposition for translocations and the resultant fusion genes and that, in turn, these fusion genes create unique transcriptional programs that promote angiogenic phenotypes in these p53-deficient backgrounds.

Specifically, kinase fusion genes involving *FLT1, PTK2,* and *AKT3* can activate key convergent gene pathways associated with blood vessel formation and remodeling (58), and they represent potential therapeutic targets for kinase inhibitors (35,59). When we consider that sarcomas have the highest frequency of kinase fusions in TCGA datasets (35), but that they also have extremely low recurrence, a more rational approach might be to develop agents that target these convergent angiogenic pathways instead of the products from the individual fusion genes themselves (60).

The two fusion genes that we confirmed by genomic structural evaluation in canine HSAs were present in approximately 20 - 40% of cells in the tumor, both genes showing chaotic amplification. Several explanations could account for these observations. One is that histology and bioinformatics assays overestimated tumor content and tumor purity. Another is that fusions are epiphenomena arising from chaotic genomes with no influence on selection. A third, which we believe is most likely, is that translocation and the resulting fusion events occur stochastically in genomically unstable cells late in the course of tumor evolution. However, the enrichment of fusion genes and angiogenic transcriptional programs suggests that these traits endow tumor cells with selective growth and/or survival advantages that contribute to tumor progression by promoting proangiogenic environments. It is worth noting that the selective pressures in the AS milieu favor not only fusion-positive clones, but also fusion-negative clones, as the establishment of a proangiogenic environment could improve survival of all the subpopulations within the tumor. Indeed, a similar mechanism might be operative in alveolar rhabdomyosarcomas, where *PAX3-FOXO1A* fusions are necessary for tumor initiation but have no effect on tumor recurrence (61,62). Further work will be necessary to distinguish which among these non-mutually exclusive possibilities are operative, and to better understand the role of fusion genes in tumor evolution of human AS and canine HSA and, potentially, in promoting clonal heterogeneity through the creation of a permissive niche.

Fusion genes have been reported in human ASs (10,63-65). For instance, one study found a *CIC-LEUTX* fusion in one of 120 (0.8%) FFPE ASs examined (10); another found a *CEP85L-ROS1* fusion in one of 34 (3.0%) ASs examined (63); and a third found an *EWSR1-ATF1* fusion in one case of AS (65). A *NUP160-SLC43A3* fusion has also been reported in the ISO-HAS AS cell line (64). However, none of these fusion genes has been identified recurrently in subsequent studies of AS samples. While these observations are consistent with our stochastic hypothesis, we cannot completely exclude the possibility that fusion genes in human ASs, or for that matter in canine HSAs, are non-pathogenic passenger aberrations.

A larger case series will be required to define the fusion gene landscape in human AS, but canine HSA provides potential insights for what might be expected. Mutations of *TP53* are largely mutually exclusive of mutations in *KDR, PIK3CA,* and *RAS* gene family in human AS, and the mutational patterns seem to be associated with the location of the primary tumor (10,12,17,66,67). We see a similar pattern emerge in a subset of canine HSA, and from our data we propose a model that can be used as a foundation to test mechanistic links between the mutational and transcriptional landscapes in malignant vascular tumors (**Fig. 4**) and determine their roles in tumor progression. In this model, inflammatory HSAs harbor no mutations of *PIK3CA,* and only rarely of *TP53,* maintaining sufficient genomic stability that disfavors formation of fusion genes. Furthermore, the transcriptional programs in these inflammatory

HSAs are weakly angiogenic, and their permissive inflammatory environments restrain growth and metastasis. Conversely, angiogenic HSAs harbor frequent mutations of *PIK3CA* and *TP53,* and fusion events. Mutations of *PIK3CA* in p53-proficient backgrounds promote pro-angiogenic environments, while in p53-deficient backgrounds, these mutations promote altered chromatin regulation and immunomodulatory transcriptional programs. Finally, mutations of *TP53* enable genomic instability with formation of fusion genes. These events are stochastic, but fusion genes that promote pro-angiogenic transcriptional programs can enhance or even supplant the effects of *PIK3CA* mutations and create environments that accelerate tumor growth and metastatic propensity.

Molecular distinctions among human ASs could be driven by their clinical phenotype and potential therapeutic responses (9,68,69). For instance, a subset of ASs harbor gene amplifications of *MYC* and *FLT4* which frequently co-occur in tumors associated with ultraviolet (UV) irradiation- or therapeutic radiation. Mutational signatures associated with UV exposure and high mutational burden might predict more favorable immunotherapeutic responses in AS patients, as they do in patients diagnosed with malignant melanoma; however, supportive clinical trials to test this premise are limited, and the use of immune checkpoint inhibitors in human AS patients thus far has yielded mixed results (70,71). The mutational signatures in canine HSA are largely confined to the “aging” (cellular replication) signature (17), and total mutational burden is relatively low (17,72), so this condition is unlikely to provide a model to address the utility of immunotherapy in this context. However, canine HSA could provide a suitable model to address other treatments, whether pharmacologic or immunologic, directed at the molecular programs that drive progression and maintenance of the tumors in both species (33). Such validation studies could alter the paradigms for diagnosis and treatment of human AS and canine HSA, as well as of other aggressive, genomically complex sarcomas that affect humans and dogs alike.

## Data availability

RNA-Seq gene expression data generated from human sarcomas are available from the TCGA Research Network (https://www.cancer.gov/tcga). Exome sequencing data from human angiosarcomas are available from The Angiosarcoma Project (https://ascproject.org), a project of Count Me In (https://joincountmein.org/). RNA-Seq data from human AS tissues are available through the Gene Expression Omnibus (GEO; http://www.ncbi.nlm.nih.gov/geo; accession number GSE163359). RNA-Seq data from canine HSA tissues are published (17,21,32,33) and available through the GEO (accession number GSE95183) and the NCBI Sequence Read Archive (accession number PRJNA562916). All other data generated from this study are available upon request to the corresponding author.

## Acknowledgements

The authors would like to acknowledge Dr. Corrie Painter for reviewing the manuscript and providing feedback. The authors acknowledge Mitzi Lewellen for assistance with inventory, database management, and editorial assistance. The authors would also like to thank Lauren Mills for processing of the next generation sequencing data and Dr. Douglas Yee, Director of Masonic Cancer Center, for assisting with the collection of human tissues. Human biospecimens were obtained from the UMN BioNet and from the CHTN. Tissue samples were provided by the CHTN which is funded by the National Cancer Institute (NCI). Other investigators may have received specimens form the same subjects. This work was partially supported by grants 1R03CA191713-01 (J.F. Modiano, A.L. Sarver, J.H. Kim) and R37CA218570 (E.K. Karlsson) from the NCI of the National Institutes of Health (NIH), grants #422 (J.F. Modiano) and 1889-G (J.F. Modiano, M. Breen, K. Lindblad-Toh) from the AKC Canine Health Foundation, grant JHK15MN-004 (J.H. Kim) from the National Canine Cancer Foundation, grant D10-501 (J.F. Modiano, M. Breen, K. Lindblad-Toh) from Morris Animal Foundation, and a grant from Swedish Cancerfonden (K. Lindblad-Toh). This work was also supported by an NIH NCI R50 grant, CA211249 (A.L. Sarver). The NIH Comprehensive Cancer Center Support Grant to the Masonic Cancer Center, University of Minnesota (P30 CA077598) provided support for the cytogenetic analyses performed in the Cytogenomics Shared Resource. K. Megquier is supported by the NCI of the NIH under Award Number F32CA247088. The content is solely the responsibility of the authors and does not necessarily represent the official views of the NIH. M. Breen is supported in part by the Oscar J. Fletcher Distinguished Professorship in Comparative Oncology Genetics at North Carolina State University. K. Lindblad-Toh is supported by a Distinguished Professor award from the Swedish Research Council. J.F. Modiano is supported by the Alvin and June Perlman Chair in Animal Oncology. The UMGC (http://genomics.umn.edu) supported for generation of genomic sequencing data libraries, and the Minnesota Supercomputing Institute (MSI) at the University of Minnesota (http://www.msi.umn.edu) provided computational resources that contributed to the results in this study. The authors gratefully acknowledge donations to the Animal Cancer Care and Research Program of the University of Minnesota that helped support this project.

## Authors’ Contributions

**Conception and design**: J.H. Kim, J.F. Modiano

**Development of methodology**: J.H. Kim, K. Megquier, A.L. Sarver, R. Thomas, J.F. Modiano **Acquisition of data (provided animals, acquired and managed patients, provided facilities, etc.)**: J.H. Kim, K. Megquier, R. Thomas, A.L. Sarver, N. Cheng, M.A. Linden, P. Murugan, L. Oseth, C.L. Foster

**Analysis and interpretation of data (e.g., statistical analysis, biostatistics, computational analysis)**: J.H. Kim, K. Megquier, R. Thomas, A.L. Sarver, J.M. Song, Y.T. Kim, N. Cheng, M.A. Linden, P. Murugan, I. Elvers, R. Swofford, J. Turner-Maier, E.K. Karlsson, M. Breen, K. Lindblad-Toh, J.F. Modiano

**Writing, review, and/or revision of the manuscript**: J.H. Kim, K. Megquier, R. Thomas, A.J. Graef, K. Lindblad-Toh, J.F. Modiano with help from all authors

**Administrative, technical, or material support**: A.J. Graef

**Study supervision**: J.H. Kim, M. Breen, K. Lindblad-Toh, J.F. Modiano

