## Supplementary Figures for "Genomically Complex Human Angiosarcoma and Canine Hemangiosarcoma Establish Convergent Angiogenic Transcriptional Programs Driven by Novel Gene Fusions"

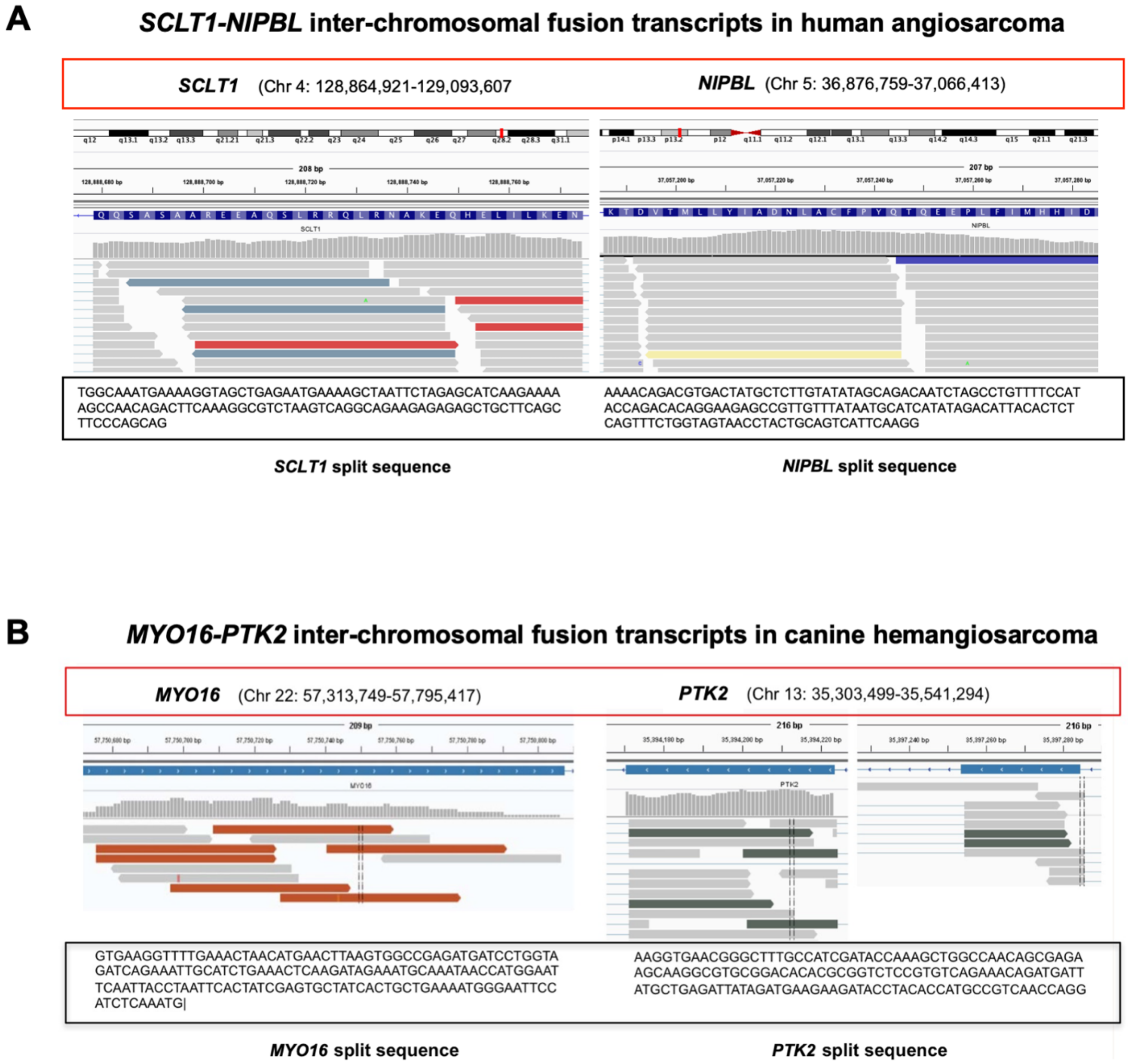


**Supplementary Fig. S1.** Representative putative inter-chromosomal fusion transcripts in human AS and canine HSA. *SCLT1-NIPBL* fusion in human angiosarcoma (**A**) and *MYO16-PTK2* fusion transcripts in canine HSA (**B**) are visualized identified by the Integrative Genomics Viewer. Mate sequenced reads are indicated in different colors.

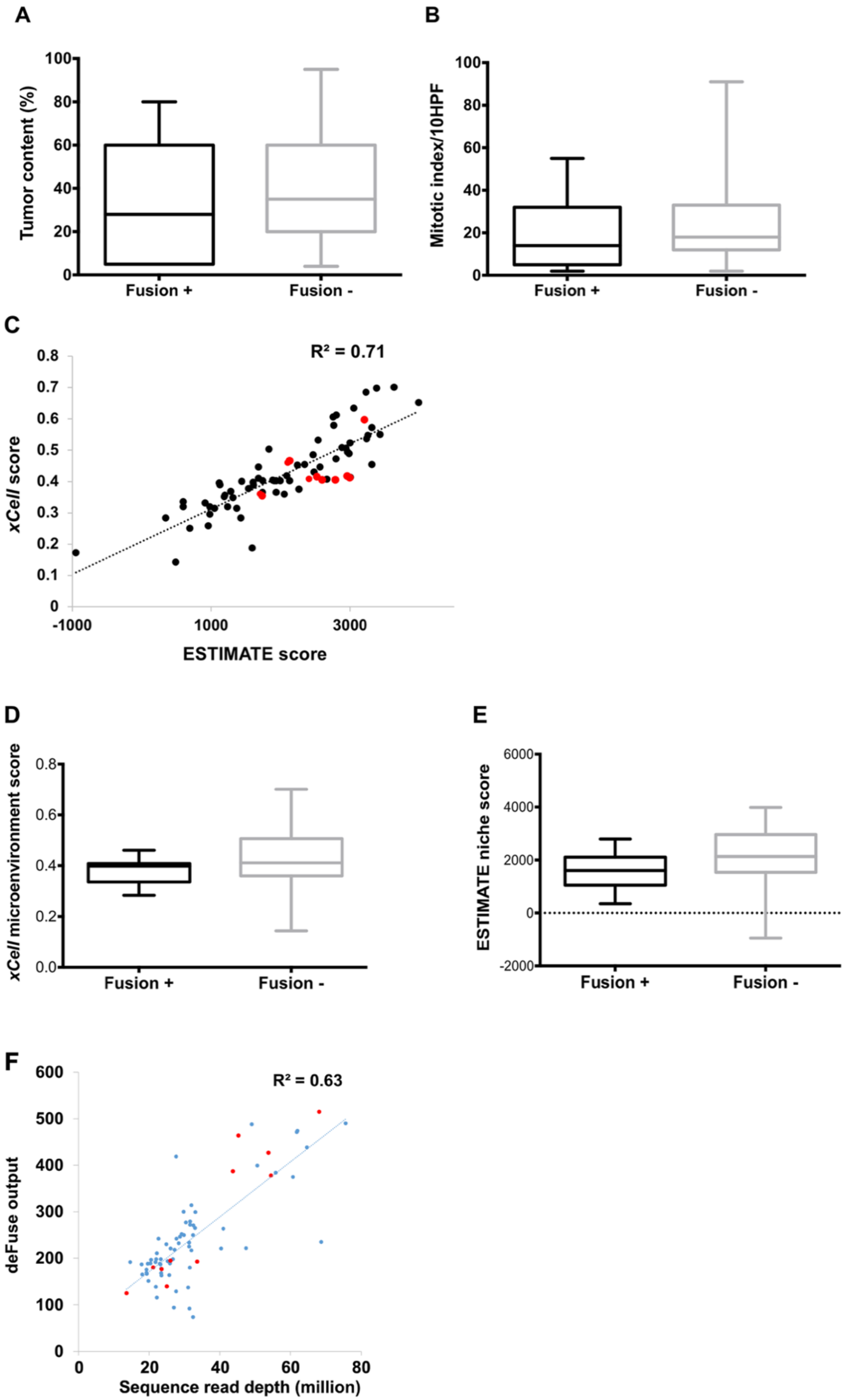
A
Tumor content (%)
100
80
60
40
20
0
Fusion +
Fusion -
B
Mitotic index/10HPF
100
80
60
40
20
0
Fusion +
Fusion -
C
R² = 0.71
xCell score
0.8
0.7
0.6
0.5
0.4
0.3
0.2
0.1
0
-1000
1000
3000
ESTIMATE score
D
xCell microenvironment score
0.8
0.6
0.4
0.2
0.0
Fusion +
Fusion -
E
ESTIMATE niche score
6000
4000
2000
0
-2000
Fusion +
Fusion -
F
R² = 0.63
deFuse output
600
500
400
300
200
100
0
0
20
40
60
80
Sequence read depth (million)

**Supplementary Fig. S2.** Fusion genes and their association with tumor content and sequencing depth in canine HSA tissues. The presence of fusion genes and their association with tumor content was evaluated based on histological tumor content (**A**) and mitotic index (**B**). Histological assessment was done in tumors with fusion genes (*n* = 11) or without fusions (*n* = 59). **C,** Bioinformatic estimation of tumor content (*n* = 78). Dot plot shows data from two independent tools to predict tumor and stromal content. Red dots indicate samples with fusion genes. **D** and **E,** Bar graphs present data of tumor content bioinformatically quantified by *xCell* (D) and ESTIMATE (E) in RNA-Seq data generated from HSAs with fusion genes (*n* = 11) or without fusions (*n* = 67). **F**, Dot plot shows the relationship between the output of the fusion gene detection algorithm (deFuse) and total sequencing read depth. $R^2$ = Coefficient of determination. Pearson's Correlation.

**A** **Human angiosarcoma**

| Fusion gene | Gene | Primer | |
|---|---|---|---|
| *SCLT1-NIPBL* | *SCLT1* | Forward | AGCATCAAGAAAAAGCCAACA |
| | *NIPBL* | Reverse | TAAACAACGGCTCTTCCTGTG |

**B**

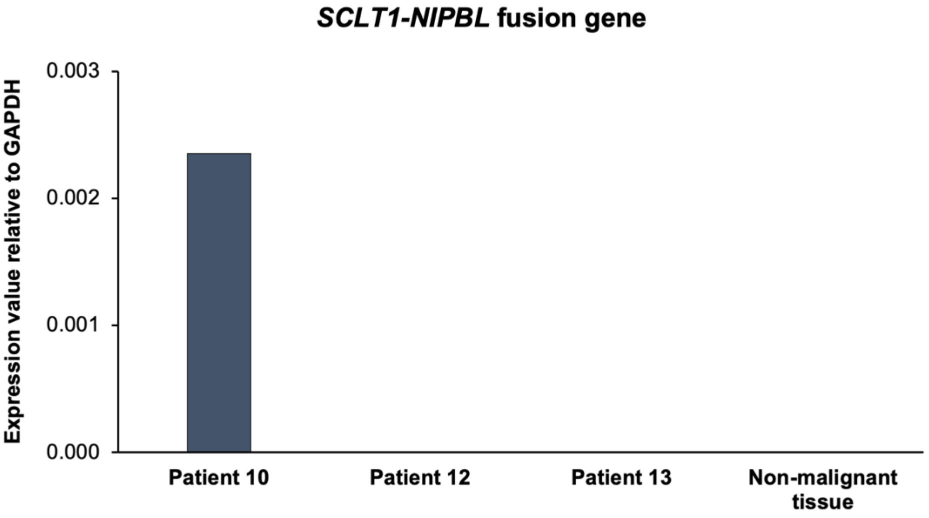


**C** **Canine hemangiosarcoma**

| Fusion gene | Gene | Primer | |
|---|---|---|---|
| *MYO16-PTK2* | *MYO16* | Forward | GCCGAGATGATCCTGGTAGA |
| | *PTK2* | Reverse | CTGTTGGCCAGCTTTGGTAT |
| *AP4E1-BAIAP2* | *AP4E1* | Forward | GCGTCAAGGCTTCTTTCTCTT |
| | *BAIAP2* | Reverse | CCAGGGCCTTCTCATAGTTCT |
| *AKT3-XPNPEP1* | *AKT3* | Forward | GCTACCGCCTGAATAGCTTCT |
| | *XPNPEP1* | Reverse | AATGGCAGCCTCCAGGAA |
| *NOL10-PTPRB* | *NOL10* | Forward | GCAGCAGACAGAGTCAGCAC |
| | *PTPRB* | Reverse | TGTGCTTTCCGGATTCTCTT |

**D**

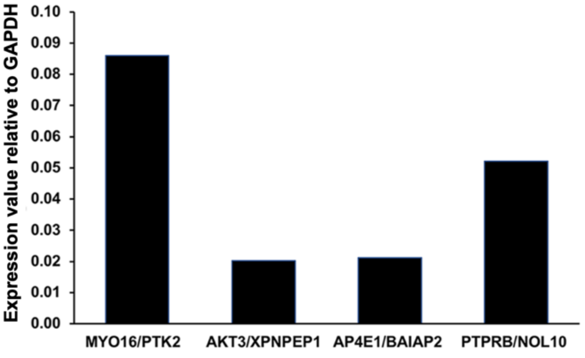


**E**

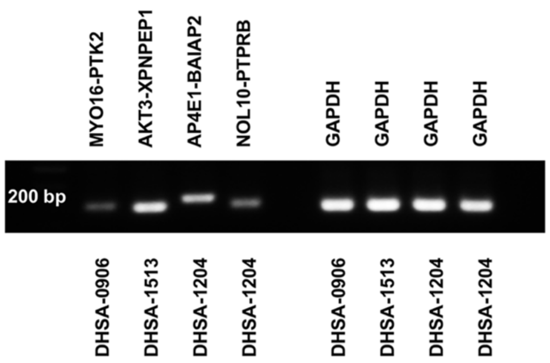

**Supplementary Fig. S3. A**, Primers designed for amplification of *SCLT1-NIPBL* fusion transcripts identified in human angiosarcoma. **B**, *SCLT1-NIPBL* fusion gene amplification by quantitative RT-PCR. Expression value represents $2^{(-\Delta Ct)(\text{Fusion gene/GAPDH})}$. **C**, Primers designed for amplification of four fusion transcripts identified in canine hemangiosarcomas. **D**, Fusion gene amplification was confirmed in canine tumors by quantitative RT-PCR. **E**, Electrophoresis (1% agarose gel) of DNA product by RT-PCR was done.

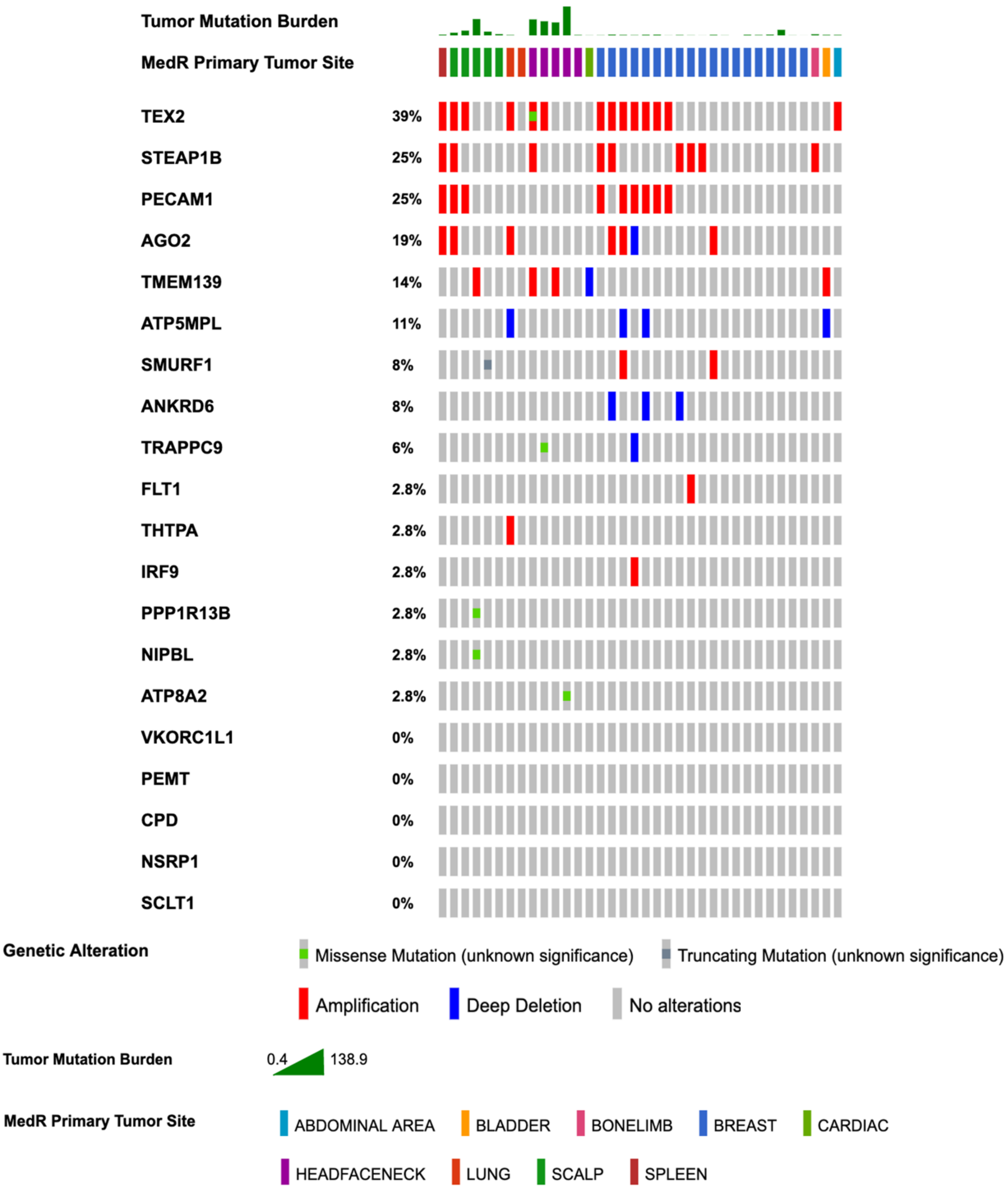
Tumor Mutation Burden
MedR Primary Tumor Site
TEX2 39%
STEAP1B 25%
PECAM1 25%
AGO2 19%
TMEM139 14%
ATP5MPL 11%
SMURF1 8%
ANKRD6 8%
TRAPPC9 6%
FLT1 2.8%
THTPA 2.8%
IRF9 2.8%
PPP1R13B 2.8%
NIPBL 2.8%
ATP8A2 2.8%
VKORC1L1 0%
PEMT 0%
CPD 0%
NSRP1 0%
SCLT1 0%
Genetic Alteration
Missense Mutation (unknown significance)
Truncating Mutation (unknown significance)
Amplification
Deep Deletion
No alterations
Tumor Mutation Burden
0.4
138.9
MedR Primary Tumor Site
ABDOMINAL AREA
BLADDER
BONELIMB
BREAST
CARDIAC
HEADFACENECK
LUNG
SCALP
SPLEEN

**Supplementary Fig. S4.** Genetic alterations of fusion partner genes identified in human AS. OncoPrint was used to visualize tumor mutation burden, somatic variations, and DNA copy number alteration of 20 fusion partner genes identified in RNA-Seq data of human ASs. Data were retrieved from Exome-sequencing of 36 human ASs in cBioPortal database for Cancer Genomics (The Angiosarcoma Project - Count Me In, Ref #12).

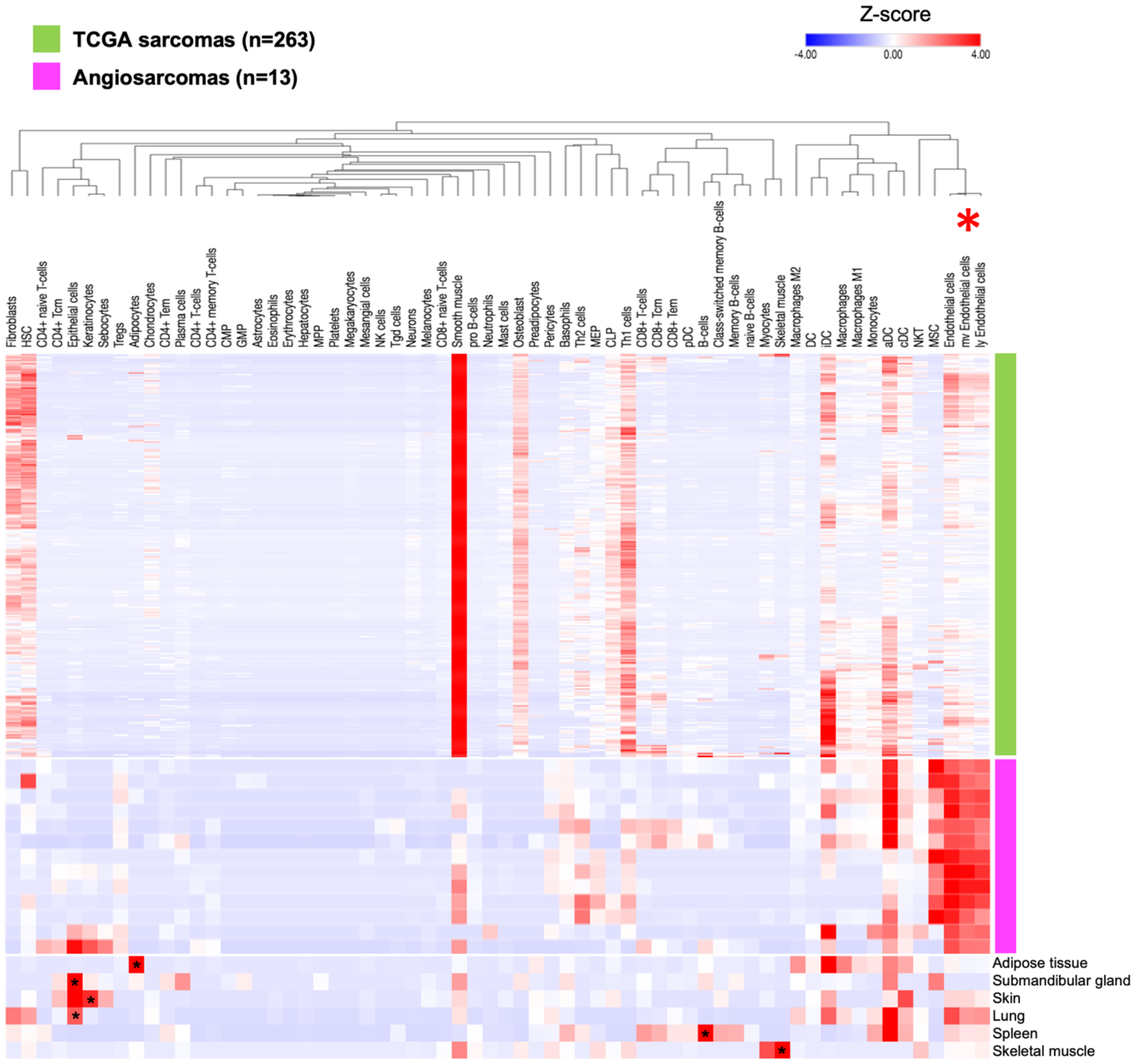


Abbreviation

| | | | |
|---|---|---|---|
| Adipocytes | Class-switched memory B-cells | Macrophages M2 | Preadipocytes |
| Astrocytes | Dendritic cell (DC) | Mast cells | Sebocytes |
| B-cells | Endothelial cells | Megakaryocytes | Skeletal muscle |
| Basophils | Eosinophils | Melanocytes | Smooth muscle |
| CD4+ T-cells | Epithelial cells | Memory B-cells | Tgd (gamma delta T) cells |
| CD4+ Tcm | Erythrocytes | Mesangial cells | Th1 (type 1 T helper) cells |
| CD4+ Tem | Fibroblasts | Monocytes | Th2 (type 2 T helper) cells |
| CD4+ memory T-cells | GMP (granulocyte macrophage progenitor) | Myocytes | Tregs (regulatory T cells) |
| CD4+ naive T-cells | HSC (Hematopoietic stem cell) | NK cells (natural killer) | Activated DC (aDC) |
| CD8+ T-cells | Hepatocytes | NKT (natural killer T) | Classical DC (cDC) |
| CD8+ Tcm (central memory) | Keratinocytes | Neurons | Immature DC (iDC) |
| CD8+ Tem (effector memory) | MEP (megakaryocytic and erythroid progenitor) | Neutrophils | Lymphatic (ly) Endothelial cells |
| CD8+ naive T-cells | MPP (multipotent progenitor) | Osteoblast | Microvascular (mv) Endothelial cells |
| CLP (common lymphoid progenitor) | MSC (mesenchymal stem cell) | Pericytes | Naive B-cells |
| CMP (common myeloid prohenitor) | Macrophages | Plasma cells | Plasmacytoid DC (pDC) |
| Chondrocytes | Macrophages M1 | Platelets | Pro B-cells |

**Supplementary Fig. S5.** Cell type enrichment analysis in ASs and TCGA sarcomas. Heatmap shows cell type enrichment analysis performed by *xCell* tool, predicting 64 different immune and stroma cell types from RNA-Seq gene expression data. Results of the analyses from TCGA sarcomas ($n$ = 263), AS samples ($n$ = 13), and non-malignant tissues ($n$ = 6) were normalized by Z-score. Red asterisk (*) indicates a gene signature of endothelial cells including microvascular and lymphatic endothelial cells. Black asterisks (*) indicate gene signatures of cell type representing the corresponding non-malignant tissues.

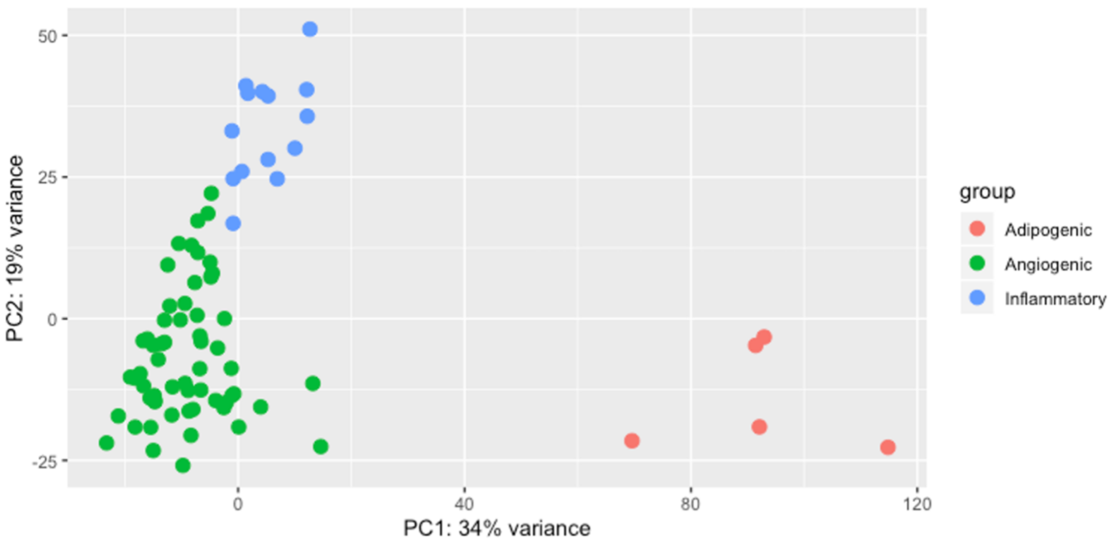


**Supplementary Fig. S6.** Three distinct molecular groups of canine HSAs. Principal Component Analysis shows distinct molecular groups of canine HSAs (*n* = 78; including two technical replicates) using 18,495 protein coding genes.

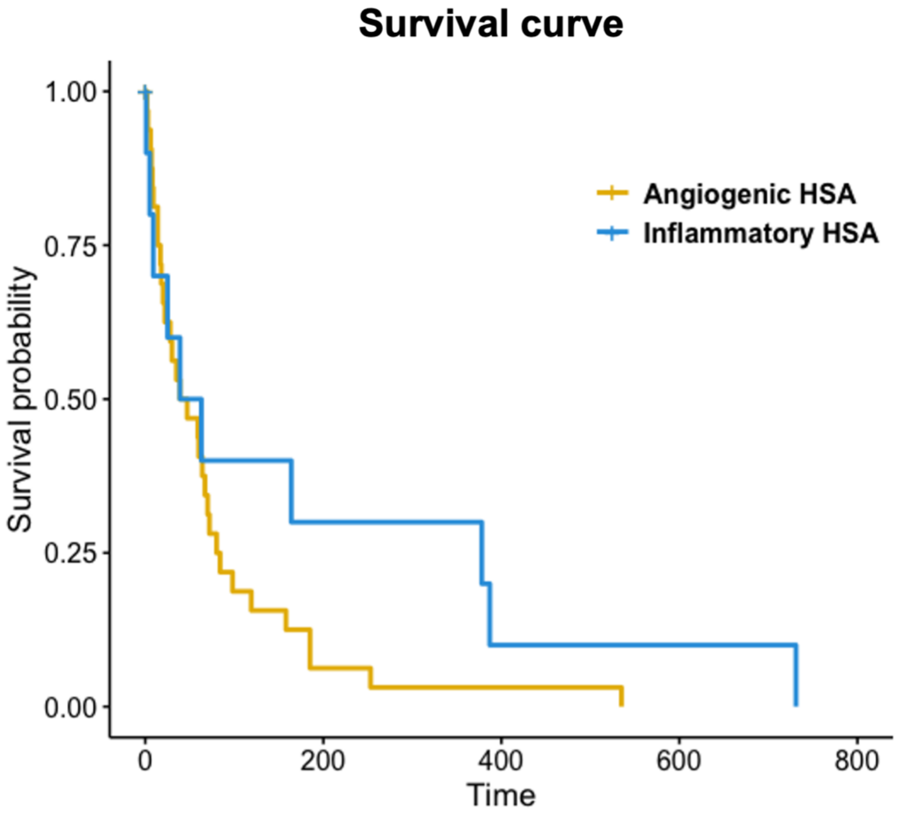


**Supplementary Fig. S7.** Kaplan-Meier probability of survival for dogs with angiogenic and inflammatory HSA. Kaplan-Meier curve shows overall survival time (days) of dogs with angiogenic HSA ($n$ = 32; yellow line) and inflammatory HSA ($n$ = 10; blue line) ($P$ = 0.19; log rank test). Dogs euthanized at diagnosis were censored.

**A**

| Age at diagnosis | Breed | Sex | Neutered | Affected organ |
|---|---|---|---|---|
| >= Median (N=39) | Golden Retriever (N=43) | Female (N=36) | Neutered (N=53) | Spleen (N=44) |
| < Median (N=32) | Labrador Retriever (N=3) | Male (N=38) | Intact (N=12) | Liver (N=9) |
| N/A (N=3) | Portuguese Water Dog (N=7) | | N/A (N=9) | Heart (N=15) |
| | Keeshond (N=2) | | | Other (N=6) |
| | German Shepherd Dog (N=6) | | | |
| | Other (N=9) | | | |
| | Mixed (N=4) | | | |

**B**

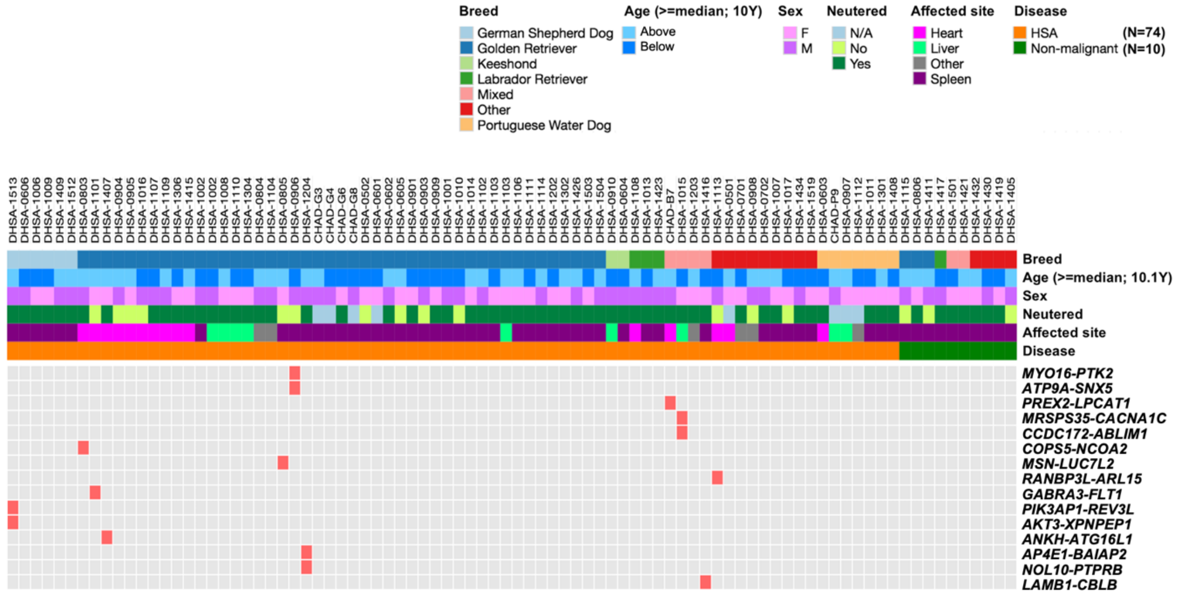


**Supplementary Fig. S8.** Association of the fusion events with demographic and histopathologic features in canine HSA. **A,** Table displays dog signalment and tumor organs obtained from dogs affected with HSA. **B**, Heatmap illustrates presence of fusion genes and the sample distribution.

A

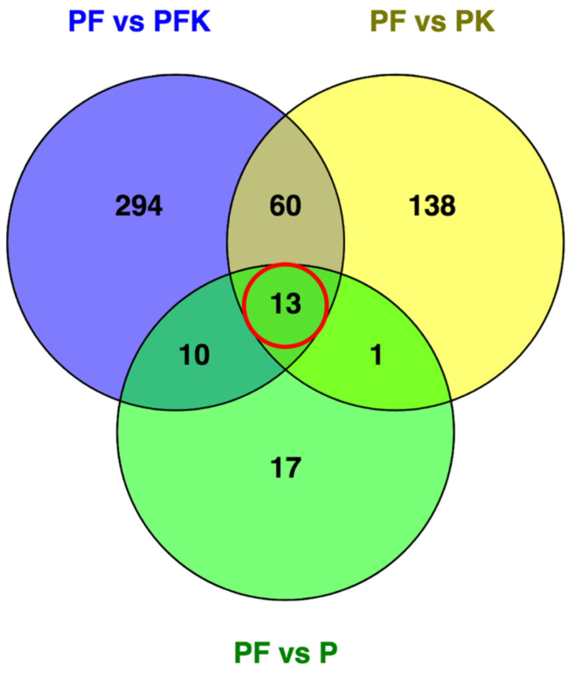


B

| Gene | P-value | Fold change | FDR p-value correction |
|---|---|---|---|
| ***TNN*** | 3.40E-17 | 113.38 | 8.24E-13 |
| ***NOX5*** | 9.37E-14 | 68.85 | 7.56E-10 |
| ***GRM3*** | 2.35E-09 | 59.29 | 7.11E-06 |
| ***LECT1*** | 4.79E-07 | 44.07 | 5.06E-04 |
| ***FMOD*** | 5.07E-08 | 39.48 | 8.19E-05 |
| ***OGN*** | 1.15E-08 | 29.61 | 2.34E-05 |
| ***COL11A1*** | 1.20E-09 | 28.08 | 4.14E-06 |
| ***CPXM2*** | 1.96E-07 | 22.85 | 2.49E-04 |
| ***C2orf40*** | 2.10E-08 | 22.41 | 3.91E-05 |
| ***COMP*** | 1.83E-14 | 19.81 | 2.22E-10 |
| ***LEPR*** | 1.51E-11 | 19.55 | 9.16E-08 |
| ***SFRP4*** | 1.39E-10 | 16.86 | 5.62E-07 |
| ***PDGFRL*** | 1.16E-08 | 11.32 | 2.34E-05 |

**Supplementary Fig. S9.** Gene pathways associated with commonly enriched genes in tumors with fusion genes and *TP53* mutations in canine HSA. **A,** Venn diagram displays numbers of differentially expressed genes and their relationship between group comparisons. **B,** Table shows a list of 13 genes that are commonly enriched in PF tumor group compared to other groups (P, PK, PFK). PF = *TP53* mutation with fusion gene; PFK = Co-mutation of *TP53* and *PIK3CA* and fusion gene; PK = Co-mutation of *TP53* and *PIK3CA* without fusion gene; P = *TP53* mutation with neither *PIK3CA* mutation nor fusion gene.

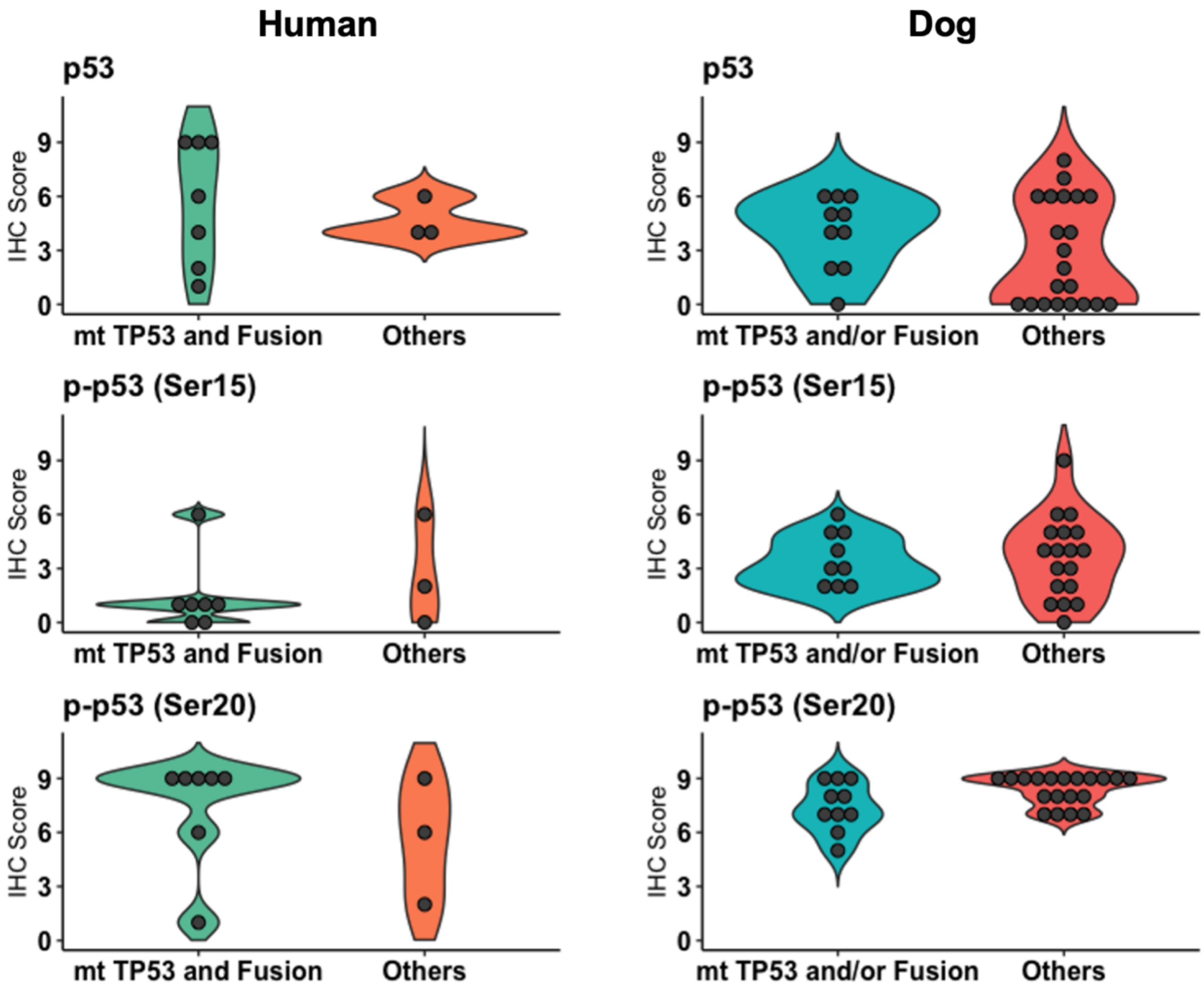


**Supplementary Fig. S10.** Violin plots display IHC scores of p53, p-p53 (Ser15), and p-p53 (Ser20) in human AS and canine HSA. Left panel: IHC scores are compared between human angiosarcomas with *TP53* mutation and fusion genes and the tumors without those. Right panel: IHC scores are compared between canine HSAs having *TP53* mutation and/or fusion genes and the tumors without any of those.

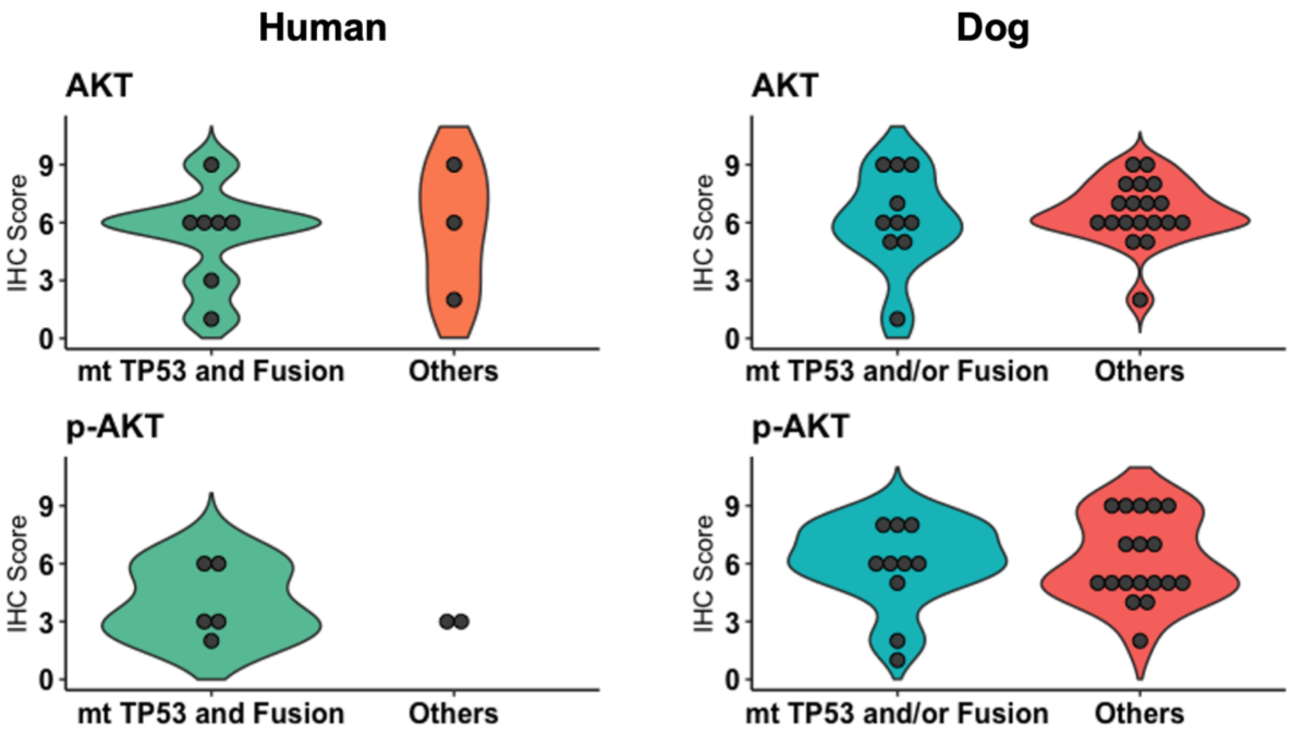


**Supplementary Fig. S11.** Violin plots present IHC scores of AKT and p-AKT (Thr308) in human AS and canine HSA. Left panel: IHC scores are compared between human angiosarcomas with *TP53* mutation and fusion genes and the tumors without those. Right panel: IHC scores are compared between canine HSAs having *TP53* mutation and/or fusion genes and the tumors without any of those.

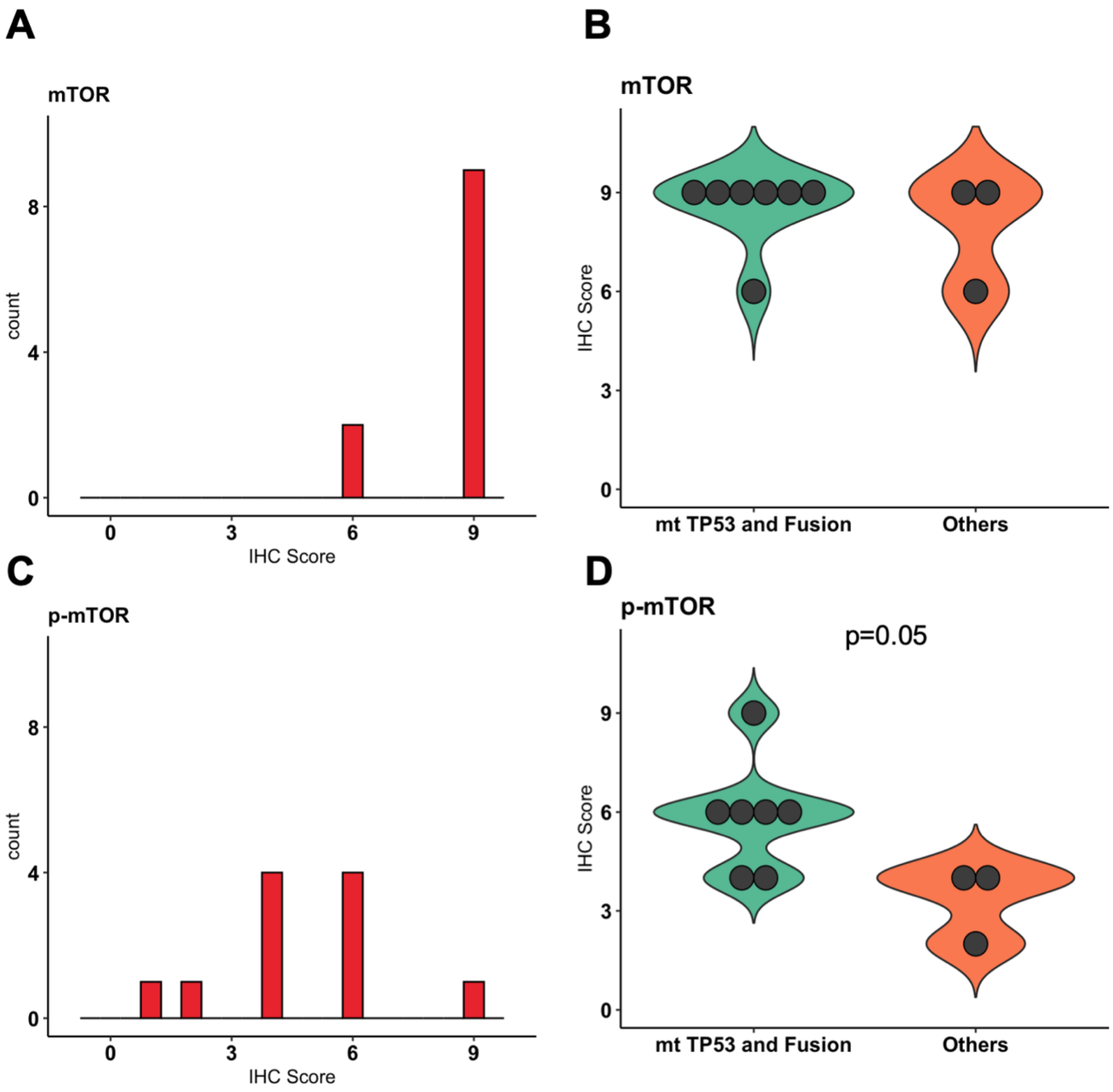


**Supplementary Fig. S12.** IHC scores of mTOR and p-mTOR (Ser2448) in human AS. IHC scores of mTOR protein are presented in bar graph (**A**) and compared between the tumors with *TP53* mutation and fusion genes and without those genetic alterations (**B**). IHC scores of p-mTOR protein are presented in bar graph (**C**) and compared with the tumors containing *TP53* mutation and fusion genes and without those genetic alterations (**D**). *P*-value is calculated in two-tailed Mann-Whitney test.
